# Bioactivity descriptors for uncharacterized compounds

**DOI:** 10.1101/2020.07.21.214197

**Authors:** Martino Bertoni, Miquel Duran-Frigola, Pau Badia-i-Mompel, Eduardo Pauls, Modesto Orozco-Ruiz, Oriol Guitart-Pla, Víctor Alcalde, Víctor M Diaz, Antoni Berenguer-Llergo, Antonio García de Herreros, Patrick Aloy

## Abstract

Chemical descriptors encode the physicochemical and structural properties of small molecules, and they are at the core of chemoinformatics. The broad release of bioactivity data has prompted enriched representations of compounds, reaching beyond chemical structures and capturing their known biological properties. Unfortunately, ‘bioactivity descriptors’ are not available for most small molecules, which limits their applicability to a few thousand well characterized compounds. Here we present a collection of deep neural networks able to infer bioactivity signatures for any compound of interest, even when little or no experimental information is available for them. Our ‘signaturizers’ relate to bioactivities of 25 different types (including target profiles, cellular response and clinical outcomes) and can be used as drop-in replacements for chemical descriptors in day-to-day chemoinformatics tasks. Indeed, we illustrate how inferred bioactivity signatures are useful to navigate the chemical space in a biologically relevant manner, unveiling higher-order organization in natural product collections, and to enrich mostly uncharacterized chemical libraries for activity against the drug-orphan target Snail1. Moreover, we implement a battery of signature-activity relationship (SigAR) models and show a substantial improvement in performance, with respect to chemistry-based classifiers, across a series of biophysics and physiology activity prediction benchmarks.

## Introduction

Most of the chemical space remains uncharted and identifying its regions of biological relevance is key to medicinal chemistry and chemical biology^1,2^. To explore and catalogue this vast space, scientists have invented a variety of chemical descriptors, which encode physicochemical and structural properties of small molecules. These encodings are at the core of chemoinformatics and are fundamental in compound similarity searches, clustering and, when applied to computational drug discovery (CDD), structure optimization and target prediction.

The corpus of bioactivity records available suggests that other numerical representations of molecules are possible, reaching beyond chemical structures and capturing their known biological properties. Indeed, it has been shown that an enriched representation of molecules can be achieved through the use of ‘bioactivity signatures’^3^. Bioactivity signatures are multi-dimensional vectors that capture the biological traits of the molecule (for example, its target profile) in a format that is akin to the structural descriptors or fingerprints used in the field of chemoinformatics. Currently, public databases contain experimentally determined bioactivity data for about a million molecules, which represent only a small percentage of commercially available compounds^4^ and a negligible fraction of synthetically accessible chemical space^5^. In practical terms, this means bioactivity signatures cannot be derived for most compounds, and CDD methods are limited to using chemical information alone as a primary input, thereby hindering their performance and not fully exploiting the bioactivity knowledge produced over the years by the scientific community.

Recently, we integrated the major chemogenomics and drug databases in a single resource named the Chemical Checker (CC), which is the largest collection of small molecule bioactivity signatures available to date^6^. In the CC, bioactivity signatures are organized by data type (ligand-receptor binding, cell sensitivity profiles, toxicology, etc.), following a chemistry-to-clinics rationale that facilitates the selection of relevant signature classes at each step of the drug discovery pipeline. In essence, the CC is an alternative representation of the small-molecule knowledge deposited in the public domain and, as such, it is also limited by the availability of experimental data and the coverage of its source databases (e.g. ChEMBL^7^ or DrugBank^8^). Thus, the CC is most useful when a substantial amount of bioactivity information is available for the molecules and remains of limited value for poorly characterized compounds^9^. In the current study, we present a methodology to *infer* CC bioactivity signatures for any compound of interest, based on the observation that the different bioactivity spaces are not completely independent, and thus similarities of a given bioactivity type (e.g. targets) can be transferred to other data kinds (e.g. therapeutic indications). Overall, we make bioactivity signatures available for any compound of interest, assigning confidence to our predictions and illustrating how they can be used to navigate the chemical space in an efficient, biologically relevant manner. Moreover, we explore their added value in the identification of hit compounds against the drug-orphan target Snail1 in a mostly uncharacterized compound library, and through the implementation of a battery of signature-activity relationship (SigAR) models to predict biophysical and physiological properties of molecules.

## Results and Discussion

The current version of the CC is organized in 5 levels of complexity (A: Chemistry, B: Targets, C: Networks, D: Cells and E: Clinics), each of which is divided into 5 sublevels (1-5). In total, the CC is composed of 25 spaces capturing the 2D/3D structures of the molecules, targets and metabolic genes, network properties of the targets, cell response profiles, drug indications and side effects, among others (Figure 1a). In the CC, each molecule is annotated with multiple n-dimensional vectors (i.e. bioactivity signatures) corresponding to the spaces where experimental information is available. As a result, chemistry (A) signatures are widely available (~10^6^ compounds), whereas cell-based assays (D) cover about 30,000 molecules and clinical (E) signatures are known for only a few thousand drugs (Figure 1b). We thus sought to infer missing signatures for any compound in the CC, based on the observation that the different bioactivity spaces are not completely independent and can be correlated.

**Figure 1.**
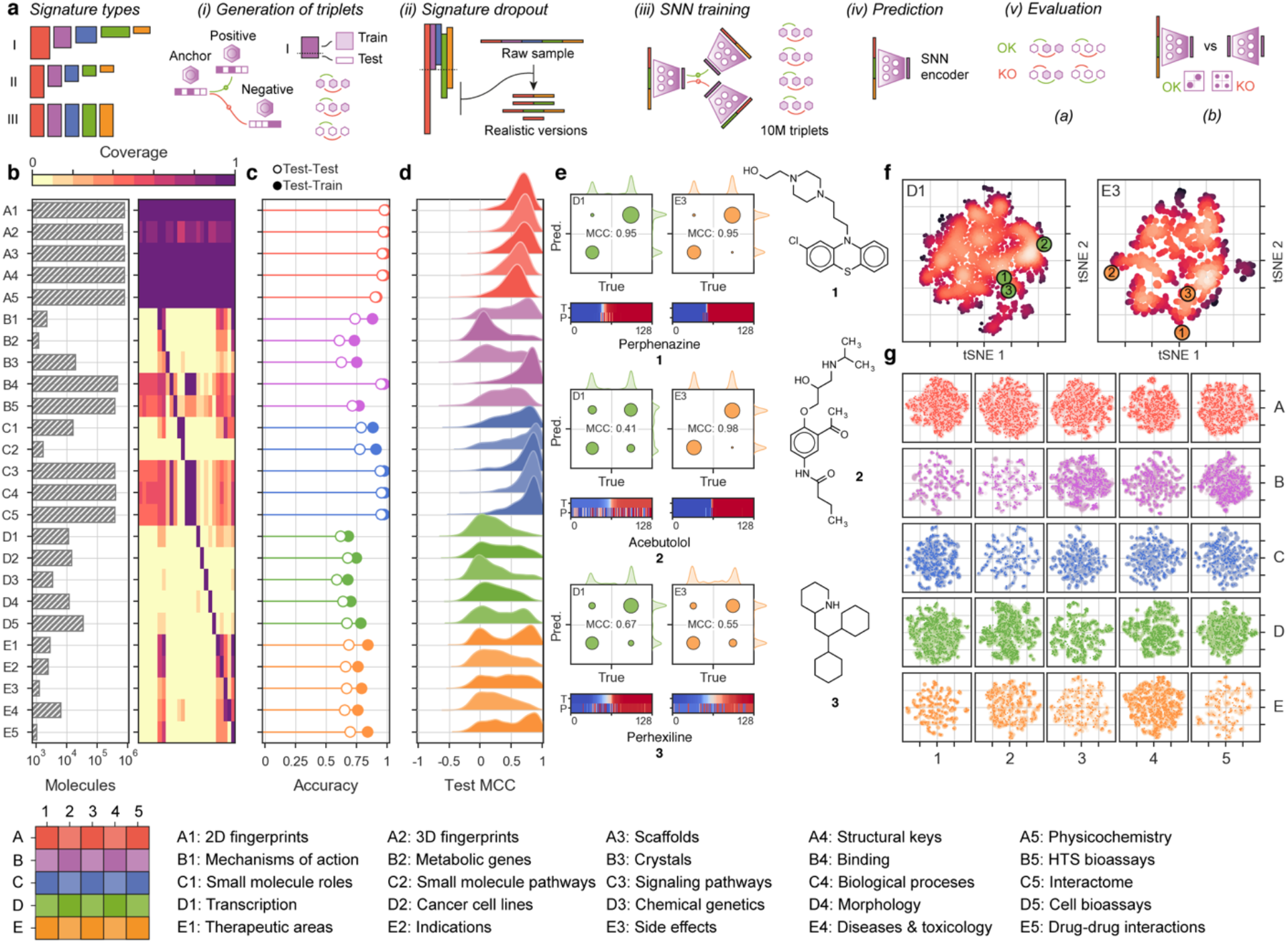
Training and evaluation of CC signaturizers. (**a**) Scheme of the methodology. Signaturizers produce bioactivity signatures that ‘fill the gaps’ in the experimental version of the CC. A SNN is trained using a signature-dropout scheme over 10^7^ triplets of molecules (anchor, positive, negative) to infer missing signatures in each bioactivity space. The inferred signatures are finally evaluated. (**b**) Coverage of the experimental version of the CC. The bar plot indicates the number of molecules available for each CC data type. The heatmap shows the cross-coverage between datasets, i.e. it is a 25×25 matrix capturing the proportion of molecules in one dataset (rows) that are also available in other datasets (columns) (**c**) Accuracy of the 25 signaturizers, measured as the proportion of correctly classified cases within a triplet. ‘Train-test’ refers to the case where the ‘anchor’ molecule belongs to the ‘test’ set, and the ‘positive’ and ‘negative’ molecules belong to the training set. ‘Test-test’ corresponds to the most difficult case where none of the three molecules within the triplet has been utilized during the training. (**d**) Performance of the 25 signaturizers, measured for each molecule as the correlation between the ‘true’ and ‘predicted’ signatures along the 128 dimensions. Given the bimodal distribution of signature values, signatures are binarized (positive/negative) and correlation is measured as a Matthew’s correlation coefficient (MCC) over the true-vs-predicted contingency table. (**e**) Three exemplary molecules (**1**, **2** and **3**) are shown for the D1 and E3 spaces. True and predicted signatures are displayed as color bars, both sorted according to true signature values. (**f**) Correspondingly, t-SNE 2D projections of D1 and E3 predictions, where **1**, **2** and **3** are highlighted. (**g**) 2D-projected train (gray) and test (colored) samples for the 25 CC spaces. The legend at the bottom specifies the A1-E5 organization of the CC.

Bioactivity signatures must be amenable to similarity calculations, ideally by conventional metrics such as cosine or Euclidean distances, so that short distances between molecule signatures reflect a similar biological behavior. Therefore, inference of bioactivity signatures can be posed as a ‘metric learning’ problem where observed compound-compound similarities of a given kind are correlated to the full repertoire of CC signatures, so that similarity measures are possible for any compound of interest, including those that are not annotated with experimental data. In practice, for each CC space (S_i_), we tackle the metric learning problem with a so-called Siamese neural network (SNN), having as input a stacked array of CC signatures available for the compound (belonging to any of the A1-E5 layers, S_1_-S_25_) and as output an n-dimensional embedding optimized to discern between similar and dissimilar molecules in S_i_. More specifically, we feed the SNN with triplets of molecules (an ‘anchor’ molecule, one that is similar to the anchor (‘positive’) and one that is not (‘negative’)), and we ask the SNN to correctly classify this pattern with a distance measurement performed in the embedding space (Figure 1a and S1). We trained 25 such SNNs, corresponding to the 25 spaces available in the CC. We used 10^7^ molecule triplets and chose an SNN embedding dimension of 128 for all CC spaces, scaling it to the norm so as to unify the distance magnitude across SNNs (see online Methods for details). As a result of this procedure, we obtained 25 SNN ‘signaturizers’ (S_1-25_), each of them devoted to one of the CC spaces (S_i_). A signaturizer takes as input the subset of CC signatures available for a molecule and produces a 128D signature that, in principle, captures the similarity profile of the molecule in the S_i_ CC space, where experimental information may not be available for the compound.

To handle the acute incompleteness of experimental signatures accessible for training the SNNs (Figure 1b), we devised a signature-dropout sampling scheme that simulates a realistic prediction scenario where, depending on the CC space of interest (S_i_), signatures from certain spaces will be available while others may not. For example, in the CC, biological pathway signatures (C3) are directly derived from binding signatures (B4), thus implying that, in a real B4 prediction case, C3 will never serve as a covariate. In practice, signature sampling probabilities for each CC space S_i_ were determined from the coverage of S_1_-S_25_ signatures of molecules *lacking* an experimental S_i_ signature. Overall, chemical information (A1-5), as well as signatures from large chemogenomics databases (e.g. B4-5), could be used throughout (Figure S2). Signatures related to the subset of drug molecules (e.g. MoA: B1, indications: E2, side-effects: E3, etc.) were mutually inclusive; however, they were more frequently dropped out in order to extend the applicability of signaturizers beyond the relatively narrow space of known drugs.

We evaluated the performance of a signaturizer S_i_ in an 80:20 train-test split both (a) as its ability to classify similar and dissimilar compound pairs within the triplets (Figures 1c and S3), and (b) as the correlation observed between each ‘predicted’ signature (i.e. obtained *without* using S_i_ as part of the input (S_1_-S_25_)) and, correspondingly, a ‘truth’ signature produced using only S_i_ (Figures 1c and S3). In the online Methods section, we further explain these two metrics, as well as the splitting and signature-dropout methods that are key to obtain valid performance estimates. In general, as expected, ‘chemistry’ (A) signaturizers performed almost perfect (Figure 1c), although these are of little added value since chemical information is always available for compounds. At the ‘targets’ levels (B), the performance of the signaturizers was high for large-scale binding data B4), while accuracy was variable at deeper annotation levels where the number of compounds available for training was smaller (e.g. MoA (B1) or for drug-metabolizing enzymes (B2)) (Figure 1d). Performance at the ‘networks’ level (C) was high, as this level is directly informed by the underlying ‘targets’ (B) level. Not surprisingly, the most challenging models were those related to cell-based (D) and clinical (E) data, probably due to the inherent complexity of these data with respect to the number of annotated molecules. On average, the accuracy of cell-based signaturizers was moderate (~0.7) and true-vs-predicted correlation of clinical signatures such as therapeutic classes (ATC; E1) was variable across molecules. The performance of SNNs varied depending on the CC space and molecule of interest, with signatures being well predicted in all spaces. Figure 1e-f illustrates this observation for three drugs (namely perphenazine (**1**), acebutolol (**2**) and perhexiline (**3**)), which have predicted signatures of variable quality in the transcriptional (D1) and side-effects (E3) spaces. Overall, bioactivity maps were well covered by test-set molecules, indicating that our SNNs are unbiased and able to generate predictions that are spread throughout the complete bioactivity landscape (Figures 1g and S4).

### Large-scale inference of bioactivity signatures

Having trained and validated the signaturizers, we massively inferred missing signatures for the ~800,000 molecules available in the CC, obtaining a complete set of 25×128-dimensional signatures for each molecule (chemicalchecker.org/downloads). To explore the reliability of the inferred signatures, we assigned an ‘applicability’ score (α) to predictions based on the following: (a) the proximity of a predicted signature to true (experimental) signatures available in the training set; (b) the robustness of the SNN output to a test-time data dropout^10^; and (c) the accuracy expected *a priori* based on the experimental CC datasets available for the molecule (Figure 2a). A deeper explanation of this score can be found in the online Methods, along with Figure S5 showing the relative contribution of a, b and c factors to the value of α. In a similarity search exercise, we found that α scores ≥ 0.5 retrieved a significant number of true hits (odds-ratios > 8, P-values < 1.7·10^−21^ (Figure S6)). This observation shows that, even for modest-quality CC spaces such as D1 (transcription), the number of signatures available can be substantially increased by our method (in this case from 11,638 molecules covered in the experimental version of the CC to 69,532 (498% increase) when SNN predictions are included (Figure S7)). Moreover, low- and high-α areas of the signature landscape can be easily delimited, indicating the presence of reliable regions in the prediction space (Figure 2b).

**Figure 2.**
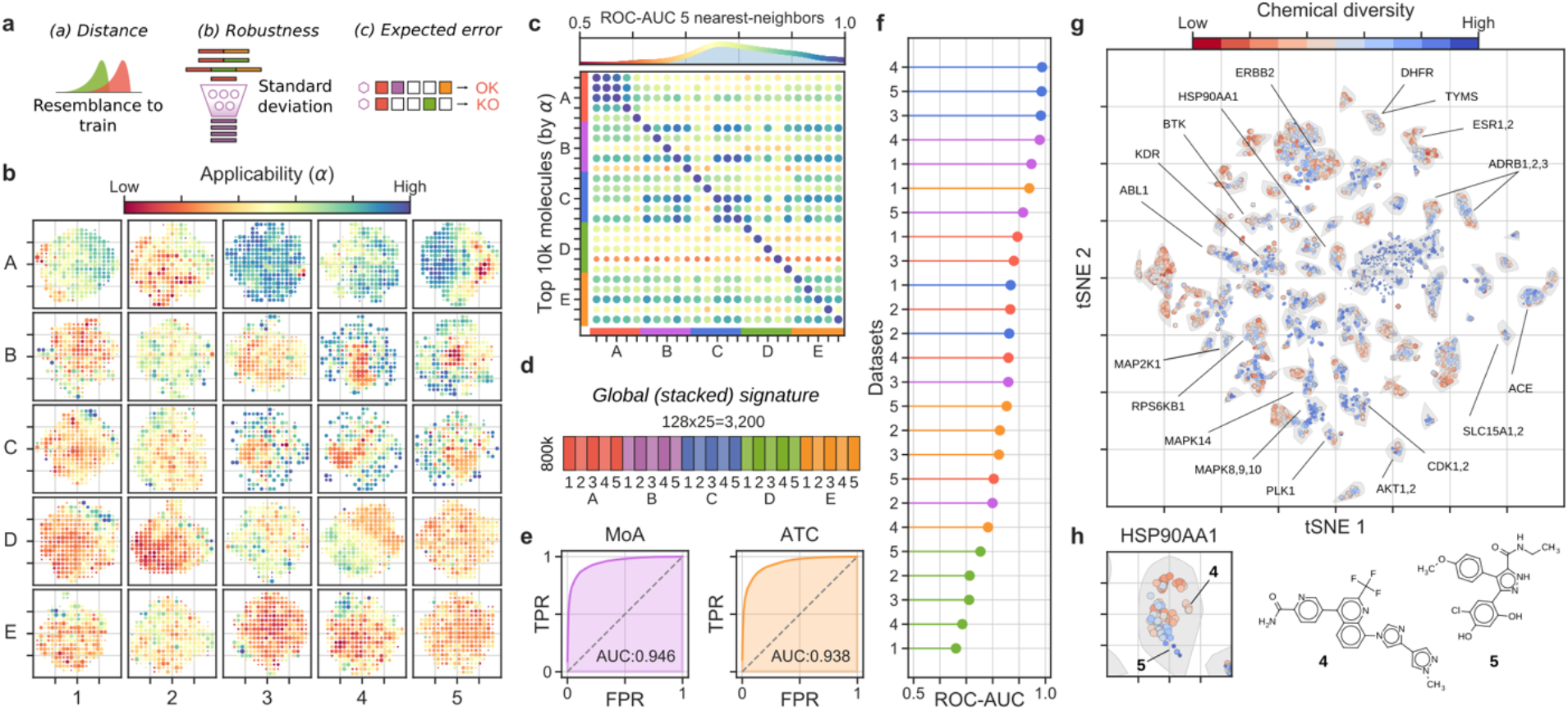
Large-scale bioactivity prediction using the signaturizers (~800k molecules). (**a**) Features combined to derive the applicability scores (α). (**b**) Applicability scores for the predictions, displayed across the 25 (A1-E5) 2D-projected signature maps. A grid was defined on the 2D coordinates, molecules were binned and the average α is plotted in a red (low) to blue (high) color scale. (**c**) Cross-correlation between CC spaces, defined as the capacity of similarities measured in S_i_ (rows) to recall the top-5 nearest neighbors in S_j_ (columns) (ROC-AUC). Top 10k molecules (sorted by α) were chosen as S_i_. (**d**) Scheme of the signature stacking procedure. Signatures can be stacked horizontally to obtain a global signature (GSig) of 3,200 dimensions. (**e**) Ability of similarity measures performed in the GSig space to identify pairs of molecules sharing the MoA (left) or ATC code (right) (ROC-AUC). (**f**) Likewise, ability of GSigs to identify the nearest neighbors found in the experimental (original) versions of the A1-E5 datasets. (**g**) t-SNE 2D projection of GSigs. The 10k molecules with the highest average α across the 25 signatures are displayed. The cool-warm color scale represents ‘chemical diversity’, red meaning that molecules in the neighborhood are structurally similar. A subset of representative clusters is annotated with enriched binding activities. (**h**) Example of a cluster enriched in heat shock protein 90 (HSP90AA1) with highlighted representative molecules with distinct (**4**) or chemically related (**5**) neighbors.

The 5×5 organization of the CC (A1-E5) was designed to capture distinct aspects of the chemistry and biology of compounds, and a systematic assessment of the original (experimental) resource revealed partial correlations between the 25 data types^6^. The original pattern of correlations was preserved among inferred signatures, especially for the high-α ones (Figures 2c and S8), thereby suggesting that the data integration performed by the SNNs conserves the genuine information contained within each data type, and implying that signatures can be stacked to provide non-redundant, information-rich representations of the molecules. For example, the 25 CC spaces can be concatenated horizontally to obtain a global signature (GSig) of 3,200 dimensions (25×128D), encapsulating in a unique signature all the bioactivities assigned to a molecule (Figure 2d). Similarity measures performed in the GSig space up-rank pairs of compounds with the same MoA or ATC code (Figure 2e) and have an overall correlation with the rest of experimental data available from the CC, capturing not only chemical similarities between molecules but also common target profiles, clinical characteristics and, to a lesser degree, cell-based assay read-outs (Figure 2f).

Indeed, as shown in Figure 2g, a 2D projection of GSigs reveals clusters of molecules with specific biological traits. Of note, some of the clusters group molecules with similar chemistries (e.g. ESR1,2 ligands), while others correspond to sets of diverse compounds (e.g. MAPK8,9,10 inhibitors). Most of the clusters have a mixed composition, containing subgroups of chemically related compounds while also including distinct molecules, as is the case for the HSP90AA1-associated cluster, of which compounds **4** and **5** are good representatives (Figure 2h).

### Bioactivity-guided navigation of the chemical space

Taken together, CC signatures offer a novel bioactivity-driven means to organize chemical space, with the potential to unveil higher levels of organization that may not be apparent in the light of chemical information alone. In Figure 3a, we analyze a diverse set of over 30 compound collections, ranging from species-specific metabolomes to purchasable building-block (BB) libraries. To expose the regions of the global bioactivity space covered by these collections, we first performed a large-scale GSig-clustering on the full CC. We then calculated GSigs for each compound in each library and mapped them to the CC clusters, thereby obtaining a specific cluster occupancy vector for each collection. Finally, we used these vectors to hierarchically group all the compound libraries. As can be seen, drug-related libraries (e.g. IUPHAR and IDG) had similar occupancy vectors to the reference CC library, meaning they were evenly distributed in the bioactivity space, which is expected given the over-representation of medicinal chemistry in our resource. Libraries containing BBs from different providers (ChemDiv, Sigma Aldrich and ChemBridge) were grouped together, although with an uneven representativity of the CC bioactivity space. Similar trends were observed for species-specific metabolomes (Yeast, *E. coli* and Human (HMDB)) and natural products collected from various sources (Traditional Chinese Medicines (TCM), African substances (AfroDb) or food ingredients (FooDB)).

**Figure 3.**
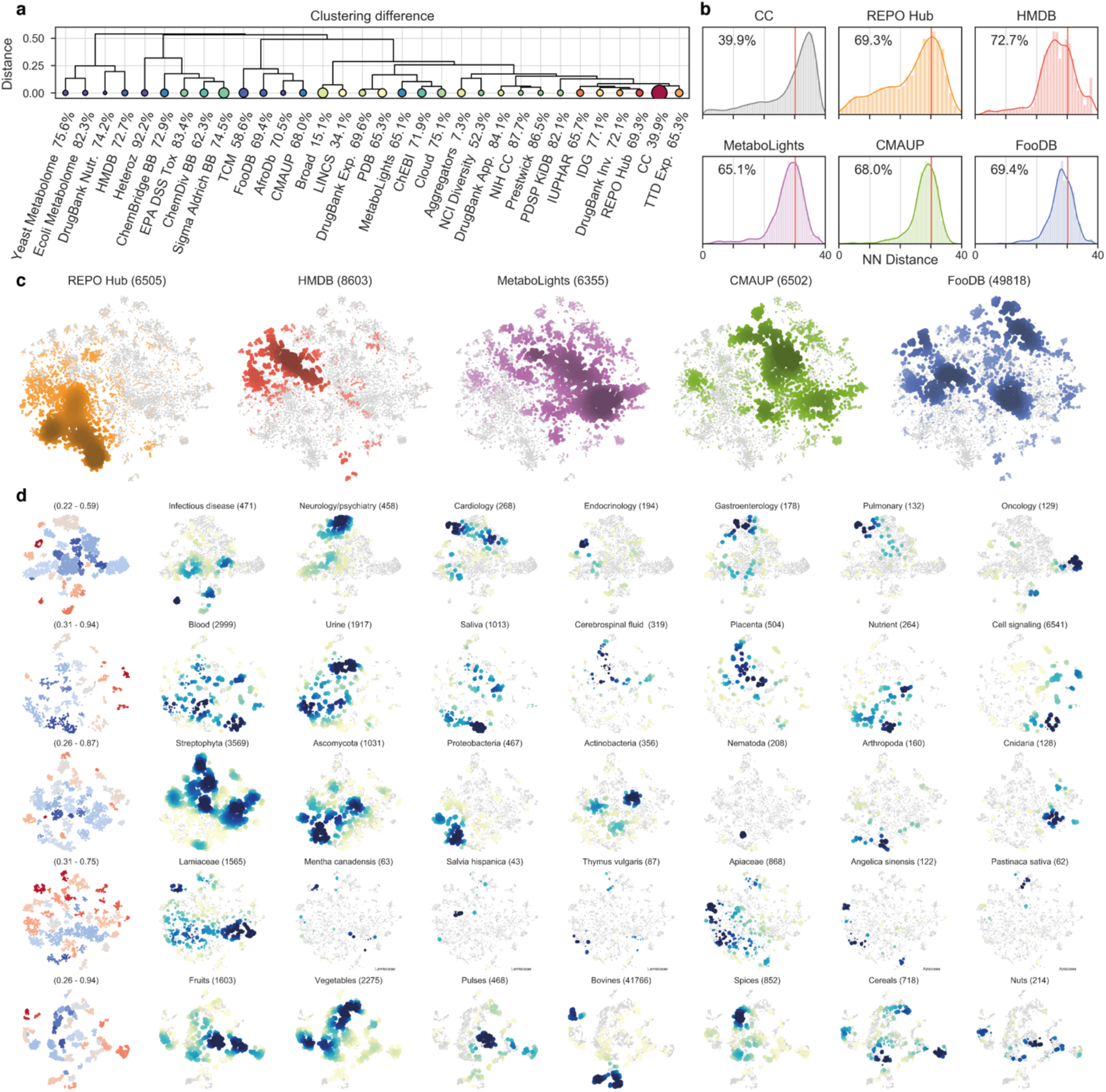
Signature-based analysis of compound collections. (**a**) Chemical libraries are hierarchically clustered by their proximity to the full CC; here, proximity is determined by the cluster occupancy vector relative to the k-means clusters identified in the CC collection (number of clusters = (N/2)^1/2^; GSigs are used). Proximal libraries have small Euclidean distances between their normalized occupancy vectors. Size of the circles is proportional to the number of molecules available in the collection. Color (blue-to-red) indicates the homogeneity (Gini coefficient) of the occupancy vectors relative to the CC. (**b**) Occupancy of high-applicability regions is further analyzed for five collections (plus the full CC). In particular, we measure the average 10-nearest-neighbor L2-distance (measured in the GSig space) of molecules to the high-α subset of CC molecules (10^3^, see Figure 1). The red line denotes the distance corresponding to an empirical similarity P-value of 0.01. The percentage indicates the number of molecules in the collection having high-α vicinities that are, on average, below the significance threshold. This percentage is shown for the rest of the libraries in panel a. (**c**) The previous five compound collections are merged and projected together (t-SNE). Each of them is highlighted in a different color. (**d**) Detail of the compound collections. The first column shows the chemical diversity of the projections, measured as the average Tanimoto similarity of the 5-nearest neighbors. Blue denotes high diversity and red high structural similarity between neighboring compounds. Coloring is done on a per-cluster basis. The rest of the columns focus on annotated subsets of molecules. Blue indicates high-density regions.

To gain a better understanding of the bioactivity areas encompassed by each collection, we chose five examples related to drug molecules, metabolomes and natural product extracts. More specifically, we considered 6,505 approved and experimental drugs (REPO Hub)^11^, 8,603 endogenous human metabolites (HMDB)^12^, 6,355 metabolites found in other species beyond vertebrates (MetaboLights)^13^, 49,818 food constituents (FooDB; www.foodb.ca) and 6,502 plant chemicals (CMAUP)^14^. Figure 3b shows that, despite their variable depth of annotation (Figure S9), these collections, for the most part, are laid out in high-α regions of the GSig space. Moreover, Figure 3c offers a comparative view of the bioactivity areas occupied by each collection, with some overlapping regions as expected, especially between natural product collections. The map reveals a region that is specific to drug molecules, possibly belonging to a set of bioactivities that is outside the reach of natural metabolites.

A deeper dive reveals further structure in the bioactivity maps. For example, when we focus on drug molecules (REPO Hub), broad therapeutic areas such as infectious diseases, neurology/psychiatry, cardiology and oncology can be circumscribed within certain regions of the GSig landscape (Figure 3d), and the same applies to finer-grained disease categories (indications) and mechanisms of action (Figure S10). Thus, the chemistry-to-clinics scope of GSigs provides a multi-level view of the chemical space, clustering compounds first on the basis of their targets and, in turn, keeping targets close in space if they belong to the same disease area. This is exemplified by PI3K, CDK and VEGFR inhibitors, which have their own well-defined clusters within the oncology region of the map, and by histamine receptor antagonists and acetylcholine receptor agonists, which are placed together in an area assigned to neurology/psychiatry (Figures 3d and S10).

Analogous observations can be made beyond the well-annotated universe of drug molecules, consistently organizing the chemical space in relevant ways. For example, the HMDB map highlights tissue- and biofluid-specific regions with varying degrees of chemical diversity (Figures 3d and S11), and the MetaboLights cross-species metabolome database is well organized by taxonomy (e.g. Chordata, Ascomycota, Actinobacteria), revealing conserved metabolite regions as well as species-specific ones (in general, we found the former to be less chemically diverse (Figure S11)). Likewise, plants can be organized in families and species by means of their ingredient signatures, as exemplified in Figure 3d for three *Lamiaceae* and two *Apiaceae* species. Finally, the map of food ingredients displays clear bioactivity clusters of food chemicals, adding to recent work suggesting that the food constituents landscape can be charted and exploited to identify links between diet and health^15^.

### Enriching chemical libraries for activity against Snail1

After seeing that inferred CC signatures are indeed useful to characterize large natural product collections, we sought to assess whether they are also advantageous in combination with more classical chemo-centric approaches. To this end, we performed a computational assessment of two chemical libraries, namely the Prestwick collection (PWCK) and the IRB Barcelona proprietary library (IRB). The IRB library contains >17k compounds, only 3% of which have reported bioactivities and are thus included in the CC. This library was originally designed to inhibit t-RNA synthetases by means of ambivalent small molecules displaying ATP-like and amino acid-like chemotypes. The PWCK library is considerably smaller (>1k compounds), and it is composed of well-annotated molecules over a wide range of activities (>99% of the molecules are present in the CC). Thus, the IRB and PWCK libraries represent two typical scenarios, the recycling of a targeted library, and the use of a small diversity-oriented compound collection, respectively.

We sought to enrich these libraries for activity against the product of SNAI1 gene, Snail1, a zinc-finger transcription factor with an essential role in the epithelial-to-mesenchymal transition (EMT)^16^. Being a transcription factor, Snail1 is almost “undruggable”^17^, and we looked for indirect strategies to inhibit its function. In a previous siRNA screening, we found that the knock-down of certain deubiquitinases (DUBs) significantly decreased Snail1 levels, suggesting that DUBs promote Snail1 stabilization and are required for its effects on EMT and cancer progression^18^.

We searched the literature for previous knowledge on DUB inhibition by small molecules^19–21^ and categorized DUBs on the basis of their performance in the siRNA-DUB/Snail1 screening assay (Data S1). We curated 45 DUB inhibitors, 6 of which were inhibitors of candidate DUBs in the siRNA-DUB/Snail1 assay. In parallel, we collected 5,540 compound-DUB interactions available in the CC corresponding to 15 of the DUBs. Overall, this search yielded a substantial pool of chemical matter related to DUB inhibition (Data S1).

In addition to DUBs, we considered other proteins with a well-established connection to Snail1 activity, including TGFBR1/2, ERK2, FBXL5/14, DDR2 and GSK3B^22^. We collected perturbational (e.g. shRNA) expression signatures for these genes, together with the signatures of prominent DUBs found in the siRNA-DUB/Snail1 screen. In total, we retrieved 95 transcriptional signatures from the L1000 Connectivity Map and 18 from the Gene Expression Omnibus (GEO)^23^ (see Data S1 for the full list of signatures). Each signature was converted to the CC D1 format. Finally, we derived networks-level (C) signatures for the previous Snail1-related genes by exploring their pathways (C3), biological processes (C4) and interactome neighborhoods (C5).

We then devised a strategy to select a few hundred compounds enriched for activity against Snail1 from the IRB and PWCK libraries (Figure 4a). On the one hand, we defined two ‘chemical queries’ to identify compounds that were either (i) chemically similar (P < 0.001) to well-curated DUB inhibitors, or (ii) similar to DUB inhibitors in a broader list (combined with binding data from chemogenomics resources; online Methods). On the other hand, we designed two ‘biological queries’ to capture connectivities between the biology of Snail1 and the bioactivity data available in the CC. In particular, we looked for (iii) compounds whose gene expression pattern might mimic Snail1-related transcriptional signatures (P < 0.001), and also (iv) compounds whose (putative) targets were functionally related to Snail1 (i.e. C3-5 similarities to TGFBR1/2, ERK2, etc., P < 0.001), applying a mild constraint based on D1.

**Figure 4.**
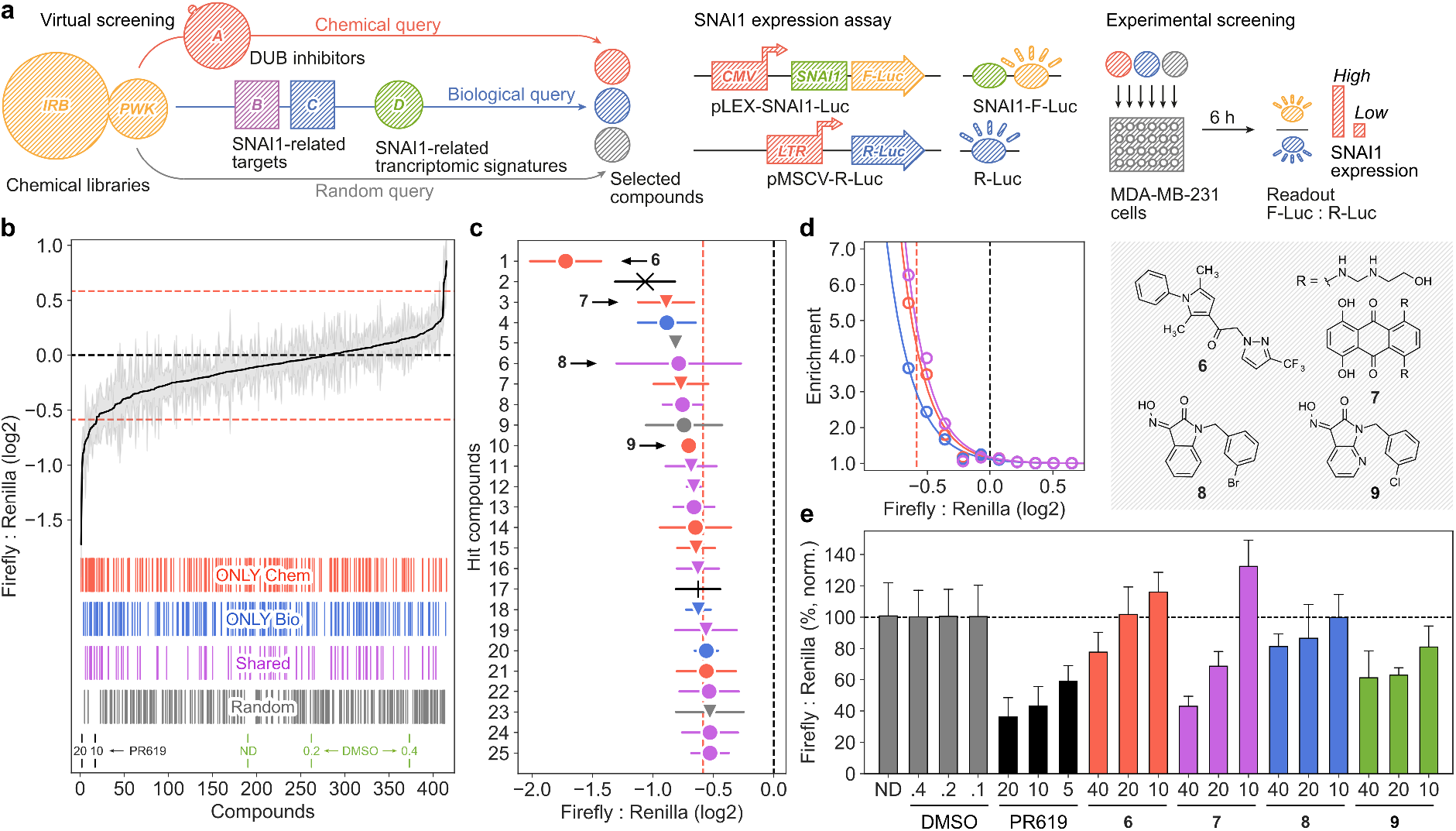
Library enrichment to identify Snail1 inhibitors. (**a**) Scheme of the methodology. Two compound libraries are screened (IRB and PWCK). A ‘chemical query’ is done by looking for similarities with known DUB inhibitors. A ‘biological query’ is done by looking for transcriptional (D1) and network-based (C3-5) signature matchings with Snail1-relevant targets. Random molecules are selected to estimate the background hit rate. A Snail1 expression assay based on Firefly:Renilla luciferase ratios is used to screen candidate compounds. (**b**) Library enrichment quantification showing the effects of compounds selected by chemical (red), biological (blue), shared between both (magenta) and random (grey) queries, as well as the positive (PR-619) and negative (DMSO) controls. (**c**) Detail of the top 25 hit compounds. (**d**) Fold enrichment of compounds selected by chemical (red), biological (blue) and shared (magenta) queries with respect to random picks, based on their capacity to modulate Snail1 levels (Firefly:Renilla assay). (**e**) MDA-MB-231 cells stably expressing luciferase constructs were treated for 6 hours with the indicated compounds, at different doses. Firefly:Renilla ratios were normalized with the corresponding concentration of vehicle (DMSO). Mean ± SD of 2 independent experiments, each of them including 4 replicas, are shown.

After inferring CC bioactivity signatures for all the ~20k compounds in our libraries, we performed the chemical and biological queries detailed above and selected 150 molecules from each query (Data S1); 183 of these belonged to the IRB library, and 117 to the PWCK collection. In addition, we selected 183 random compounds to be used as background. To validate the capacity of these compounds to decrease Snail1 protein levels, we used a Snail1-Firefly-luciferase fusion protein stably expressed in MDA-MB-231 cells (Figure 4a)^18^.

Figure 4b shows the outcome of the Snail1-luciferase screening assay. As can be seen, 22 out of the 25 compounds displaying the strongest Snail1 down-regulation (including the two controls) came from chemical and biological queries. Importantly, a substantial number of hits (10 in the top 25) were candidate molecules selected by both biological and chemical queries, and an additional 3 compounds were retrieved only by biological queries. This observation highlights the added value of bioactivity signatures to complement chemical similarity searches (Figure 4c). Overall, considering as positive those molecules able to decrease 1.5 times Snail1 levels, selected compounds showed a 6-fold enrichment over the hit-rate of random compounds (Figure 4d). It is also worth noting that 17 of the positive hits were not known to be bioactive, and therefore their CC signatures have been fully inferred by our signaturizers. Finally, we selected the 10 compounds that displayed the strongest effect on reducing Snail1 levels and re-tested them in a confirmatory dose-response assay. Indeed, 4 of them showed a dose-dependent regulation of Snail1 (Figure 4e). Of note, compounds **8** and **9** had the same chemotype, which was identified in 4 of the top 25 hits. Taken together, these results demonstrate that the various kinds of inferred chemical and biological signatures can be used to implement complex searches to tackle the activity of currently orphan targets.

### Enhanced prediction capabilities compared to chemical descriptors

In addition, we examined whether our signaturizers could be used as molecular features to predict the outcome of a given bioassay of interest, analogous to the use of chemical descriptors in structure-activity relationship (SAR) studies. We thus developed signature-activity relationship (SigAR) models, and trained machine-learning classifiers to learn discriminative features from the CC signatures of ‘active’ (1) and ‘inactive’ (0) compounds, with the goal of assigning a 1/0 label to new (untested) compounds.

To evaluate the SigAR approach in a wide range of scenarios, we used nine state-of-the-art biophysics and physiology benchmark datasets available from MoleculeNet^24^. More specifically, we considered bioassays extracted from PubChem (PCBA), namely an unbiased virtual screening dataset (MUV), inhibition of HIV replication (HIV), inhibition of beta-secretase 1 activity (BACE), blood-brain barrier penetration data (BBBP), toxicity experiments (Tox21 and ToxCast), organ-level side effects (SIDER), and clinical trial failures due to safety issues (ClinTox). Although none of these benchmark datasets are explicitly included in the CC resource, data points can be shared between MoleculeNet and the CC, which would trivialize predictions. To rule out this possibility, we excluded certain CC signature classes from some of the exercises, as detailed in Table S1 (e.g. side-effect signatures (E3) were not used in the SIDER set of MoleculeNet tasks).

Each MoleculeNet benchmark dataset has a given number of prediction tasks, ranging from 617 (ToxCast) to just one (HIV, BACE and BBBP). The number of molecules also varies (from 1,427 in SIDER to 437,929 in PCBA) (Table S1). We trained a classifier for each MoleculeNet task independently, following a conformal prediction scheme that relates the prediction score to a measure of confidence^25^. We chose to use a general-purpose machine-learning method (i.e. a random forest classifier) with automated hyperparameter tuning, allowing us to focus on the added value of the CC signatures rather than the classification algorithm. Finally, although CC signatures are abstract representations that do not offer direct structural/mechanistic interpretations, we devised a strategy to obtain high-level explanations for predicted activities. More specifically, for each molecule, we measured the cumulative explanatory potential (Shapley values^26^) of each signature type (S_1-25_) across the GSig space, indicating the classes of data (chemistry, targets, etc.) that were more determinant for the classifier decision (online Methods). In sum, we implemented an automated (parameter-free) SigAR methodology, the outcome of which can be interpreted at the signature-type level and is calibrated as a probability or confidence.

In Figures 5a-d and S12, we show the characteristics of a representative classifier, corresponding to the heat shock factor response element (SR-HSE) task in the Tox21 panel. In a 5-fold cross-validation, active molecules got higher prediction scores than inactive compounds (Figure S12). Moreover, the SigAR model outperformed the conventional chemical Morgan fingerprints (MFps) (Figure 5a). Additionally, the accuracy of the classifier was more robust to successive removal of training data (Figure 5b), suggesting that, in principle, fewer data would be necessary to achieve a proficient model if CC signatures are used. Of note, some molecules had a high prediction score with the GSig-based model but were nonetheless predicted to be inactive by the MFp-based counterpart, and *vice versa* (Figure 5c), thus pointing to complementarity between the SigAR and SAR approaches. Indeed, CC chemistry levels were *not* among the best ‘explanatory’ signature types for the SR-HSE classifier. Instead, HTS bioassays (B5) and cell morphology data (D4) appeared to be more informative (Figure 5d), an observation that is also apparent when active molecules are laid out on the B5 and D4 2D maps (Figure 5e).

**Figure 5.**
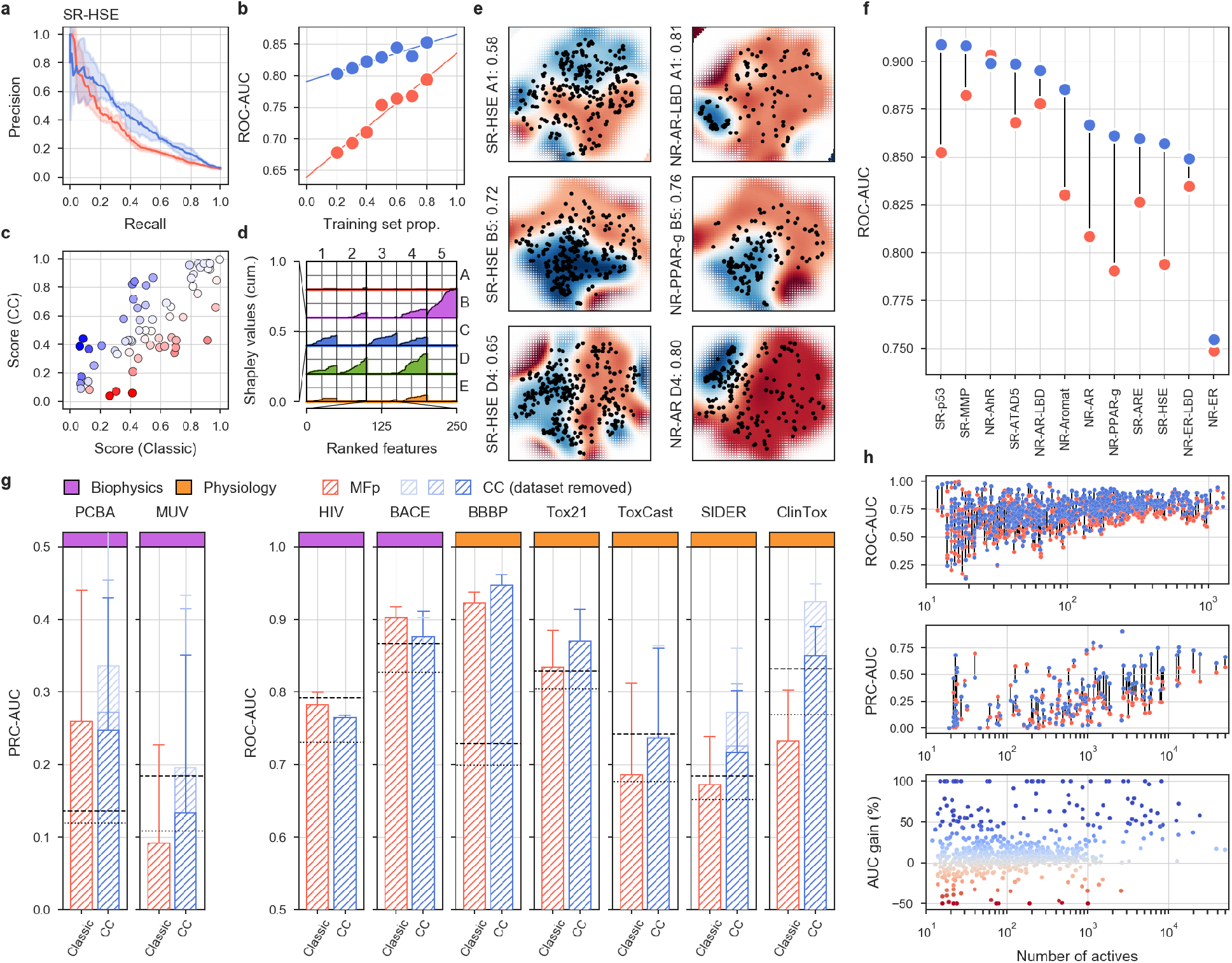
MoleculeNet benchmarks, comparing the predictive power of CC signatures with a classical MFp-based approach. (**a**) Precision-recall curves (PRCs) for the Tox21 SR-HSE task, trained with CC signatures (blue) and MFps (red). Shaded areas span the standard deviation over five stratified train-test splits. (**b**) Robustness of the SR-HSE classifier, understood as the maintenance of performance (ROC-AUC) as fewer training samples become available. (**c**) Prediction scores (probabilities) of active test molecules using MFps (x-axis) or CC signatures (y-axis). (**d**) Importance of CC datasets for the predictions. Features are ranked by their absolute Shapley value (SHAP) across samples (plots are capped at the top 250 features). For each CC dataset (S_i_), SHAPs are cumulatively summed (y-axis; normalized by the maximum cumulative sum observed across CC datasets). (**e**) 2D projections related to SR-HSE (first column) and other (second column) tasks, done for the A1, B5 and D4 CC categories (rows). A simple support vector classifier (SVC) is trained with the *(x,y)*-coordinates as features in order to determine an activity-decision function. Performance is given as a ROC-AUC on the side of the plots. Blue and red areas correspond to likely active and likely inactive regions, respectively. Active compounds are overlaid as black dots. (**f**) Performance of CC signatures (blue) and MFps (red) on the 12 Tox21 tasks. Tasks are ranked by their CC ROC-AUC performance. (**g**) Global performances of biophysics (purple) and physiology (orange) benchmark tasks. PRC and ROC AUCs are used, following MoleculeNet recommendations. Shades of blue indicate whether all 25 CC datasets were used (light) or whether conservative dataset removal was applied (darker) (Table S1). Dashed and dotted lines mark respectively the best and average reported performance in the seminal MoleculeNet study^13^. (**h**) Relative performance of CC and MFp classifiers across all MoleculeNet tasks (split by ROC-AUC and PRC-AUC metrics, correspondingly; top and middle panels). Higher performances are achieved when more active molecules are available for training (x-axis). The average gain in AUC is plotted in the bottom panel.

Figure 5f demonstrates that GSigs are generally favorable to MFps across the 12 toxicity pathways defined in the Tox21 benchmark dataset, with particularly large differences for the SR-p53, NR-Aromatase, NR-AR, NR-PPAR-gamma and SR-HSE tasks, and essentially the same performance for the NR-AhR and NR-ER tasks. Figures S13–S17 give further details for these classifiers, supporting the robustness of the SigAR approach and demonstrating that, depending on the classification task, the model will benefit from specific CC signature types (Figures 5e, S16 and S17). The NR-AhR model, for instance, mostly leverages the chemical levels (A), whereas SR-ATAD5 benefits from cell sensitivity data (D2), and NR-ER-LBD exploits the functional (e.g. biological process (C3)) information contained within the network levels of the CC.

More comprehensively, in Figure 5g we evaluate the predictive power of the SigAR classifiers across the full collection of MoleculeNet benchmark datasets, comprising 806 prediction tasks (Table S1). Our SigAR predictions were generally more accurate than the equivalent chemistry-based models, meaning that our signaturizers feed additional, valuable information to a broad range of activity prediction tasks. We observed a remarkable added value of the SigAR methodology for the physiology benchmark datasets (e.g. SIDER and ClinTox), which are, *a priori*, those that should benefit most from an integrative (data-driven) approach like ours. Overall, we observed 8.5% median improvements in performance with respect to chemistry-based classifiers (IQR: 1.4%-19.5%, Wilcoxon’s test P-value = 5·10^−60^) (Figure 5h). This implies a median reduction of the gap between actual and perfect (ideal) performance of 17.6% (IQR: 24.4%-31.5%). Reassuringly, considering only molecules with reported bioactivity (i.e. included in the CC) further accentuated the difference in performance (Figure S18), highlighting the importance of data integration methodologies to overcome the limitations of a classical (chemistry-only) approach.

### Code and models

Software for generating CC signatures is available as a python package at http://gitlabsbnb.irbbarcelona.org/packages/signaturizer. The ‘signaturizer’ API allows conversion of molecules (represented as SMILES strings) to the 25 signature types available from the CC. These pre-trained signaturizers are light-weight versions of the SNNs presented here, freeing the user from the need of setting up a full version of the CC (online Methods). Signaturizers are available as TensorFlow Hub ‘SavedModel’ instances and are automatically downloaded by the API the first time they are used. The full CC repository is open-sourced at http://gitlabsbnb.irbbarcelona.org/packages/chemical_checker.

### Concluding remarks

Drug discovery is a funneling pipeline that ends with a drug being selected from a starting pool of hundreds of thousands, if not millions, of compounds. Computational drug discovery (CDD) methods can aid in many steps of this costly process^27^, including target deconvolution, hit-to-lead optimization and anticipation of toxicity events. An efficient mathematical representation of the molecules is key to all CDD methods, 2D structural fingerprints being the default choice in many cases.

The renaissance of (deep) neural networks has fueled the development of novel structure ‘featurizers’^28^ based on graph/image convolutions of molecules^29–31^, the apprehension of the SMILES syntax^32^, or even a unified representation of protein targets^33^. These techniques are able to identify problem-specific patterns and, in general, they outperform conventional chemical fingerprints. However, neural networks remain challenging to deal with, and initiatives such as DeepChem are contributing to making them accessible to the broad CDD community^34^. The CC approach presented here shares with these initiatives the will to democratize the use of advanced molecular representations. Our approach is complementary in that it does not focus on optimally encoding chemical structures. Instead, we have undertaken the task of gathering, harmonizing and finally vectorizing the bioactivity data available for the molecules in order to embed a wide array of bioactivities in a compact descriptor.

Since CC signatures are simple 128D vectors, they are compatible with other CDD toolkits that primarily use multi-dimensional descriptors to represent molecular structures. This compatibility presents a unique opportunity to inject biological information into similarity searches, visualization of chemical spaces, and clustering and property prediction, among other widely used CDD tasks.

In this study, we showed how CC signatures can be used to navigate the chemical space in a biological-relevant manner, revealing somehow unexpected high-order structure in poorly annotated natural product collections. We also demonstrated that inferred bioactivity signatures are useful to annotate mostly uncharacterized chemical libraries and enrich compound collections for activity against a drug-orphan target, beyond chemical similarities. Moreover, compared to using chemical information alone, we observed a superior performance of SigAR models across a series of biophysics and physiology activity-prediction benchmark datasets. We chose to train models with minimal parameter tuning, illustrating how our signaturizers can be used in practice with minimal knowledge of machine learning to obtain state-of-the-art performances.

A limitation of CC signatures is that they are difficult to interpret in detail. That is, the underlying data points (binding to receptor ‘x’, occurrence of drug side effect ‘y’, etc.) cannot be deconvoluted from the 128D signature. This caveat is common to other machine-learning applications (e.g. natural language processing) where embedded representations of entities are favored over sparser, more explicit ones^35^. Nonetheless, we show that CC signatures can be interpreted at a coarser level, indicating which signature types are more informative for a certain prediction task. Another caveat of our approach is the likely existence of ‘null’ signatures corresponding to innocuous molecules with no actual bioactivity in a given CC data type^36^. Likewise, the accuracy of the signatures may vary depending on the molecule. To control for these factors, CC signatures are accompanied by an applicability score that estimates the signature quality on the basis of the amount of experimental data available for the molecule, the robustness of the prediction and the resemblance of the predicted signature to signatures available from the training set.

Contrary to most chemical descriptors, CC signatures evolve with time as bioactivity measurements accumulate in the databases. We will release updated versions of the signaturizers once a year and, as developers of the CC, we are committed to keeping abreast of the latest phenotypic screening technologies and chemogenomics datasets. Although the current version of the CC is constrained to 25 categories, our resource is prepared to accommodate new data types, offering the opportunity to customize and extend the current repertoire of signaturizers. The growth of the CC resource is restricted by the number and quality of publicly accessible datasets, a limitation that is likely to be ameliorated with the implementation of private-public partnerships and the general awareness that, in the markedly gene-centric omics era, the depth of small molecule annotation lags behind genomes and proteomes^37,38^. The ever-growing nature of chemical matter (in contrast to the finite number of genes) demands computational methods to provide a first estimate of the biological properties of compounds^39^. We believe that CC signaturizers can bridge this gap and become a reference tool to scrutinize the expected bioactivity spectrum of compounds.

## Online Methods

### Data collection

Experimental CC signatures were obtained from the CC repository (version 2019/05). Drug Repurposing Hub molecules and annotations were downloaded from https://clue.io/repurposing (June 2019). HMDB and FooDB data were downloaded from http://hmdb.ca and http://foodb.ca, respectively (April 2020). Plant ingredients were collected from CMAUP (July 2019) and cross-species metabolites from https://www.ebi.ac.uk/metabolights (April 2020). MoleculeNet benchmark datasets were downloaded from http://moleculenet.ai in June 2019. The remaining compound collections were fetched from ZINC catalogs (http://zinc.docking.org) (June 2020).

### Siamese neural networks

We carried out all procedures specified below for each CC dataset (S_i_) independently, and we trained 25 SNNs based on existing CC signatures and molecule triplets reflecting S_i_ similarities. SNNs use the same weights and neural architecture for the three input samples to produce comparable output vectors in the embedding space.

#### Covariates matrix

We trained a SNN having horizontally concatenated signatures (S_1_-S_25_) as a covariates matrix (*X*), and producing 128D-vectors as output (*Y*). The covariates matrix was stacked with a pre-compressed version of CC signatures (named signatures type II) with 128 dimensions. Only CC datasets covering at least 10% of S_i_ were stacked in *X*. Thus, given *n* molecules in S_i_, and having *m* S_1-25_ datasets cross-covering at least 10% of n, *X* would be of shape (*n*, 128·*m*). For each molecule (row), missing signatures were represented as not-a-number (NaN) values.

#### Triplet sampling

We sampled 10^7^ molecule triplets (i.e. 10^7^/n triplets per anchor molecule). Positive samples (i.e. molecules similar to the anchor) were drawn using the FAISS k-nearest neighbor search tool^40^. The value of k was empirically determined so that it maximized the average ROC-AUC of similarity measures performed against the rest of CC datasets, and it was then clipped between 10 and 50. Negative samples were randomly chosen from the pool of molecules at larger distance than the positive compounds.

#### SNN architecture

SNNs were built and trained using Keras (https://github.com/fchollet/keras). After the 128·m-dimensional input layer, we added a Gaussian dropout layer (σ = 0.1). We then sequentially added two fully connected (dense) layers whose size was determined by the m magnitude. When m·128 was higher than 512, the two hidden layers had sizes of 512 and 256, respectively. For smaller m values, we linearly interpolated the size between input and output (128) dimensions (e.g. for m = 7, the two hidden layers had sizes of 448 and 224, respectively). Finally, a dense output layer of 128 dimensions was sequentially added. For the hidden layers, we used a SeLU activation with alpha-dropout regularization (0.2), and the last (output) layer was activated with a Tanh function, together with an L2-normalization.

#### Signature dropout

We devised a dropout strategy to simulate availability of CC signatures at prediction time. To do so, we measured the proportion of experimental S_1-25_ signatures available for not-in-S_i_ molecules. These observed (realistic) probabilities were then used to mask input data at fitting time, more frequently setting those CC categories with the smaller probabilities to NaN. The S_i_ signature was dropped out with an oscillating probability (0-1) over the training iterations (5,000 oscillation cycles per epoch).

#### Loss functions

To optimize the SNN, we used a pair of loss functions with a global orthogonal regularization^41^. The first one was a conventional triplet loss, checking that the distance between the anchor and the positive molecule measured in the embedding (128D) space was shorter than the anchor-negative distance (margin = 1). The second loss was exclusively applied to the anchor molecule, and it controlled that the embedding resulting from the signature dropout was similar to the embedding obtained using S_i_ alone (mean-squared error (MSE)). Global orthogonal regularization (alpha = 1) was used to favor maximal spread-out of signatures in the embedding space. The Adam optimizer was used with a default learning rate of 10^−4^.

#### Evaluation

For each S_i_, we split the list of *n* molecules into train (80%) and test (20%) sets. Splitting was done after removing near-duplicates with FAISS. We then defined three triplet splits, i.e. train-train, test-train and test-test, using molecules from the train and test sets as anchors and positives/negatives, correspondingly. For CC spaces with less than 30,000 molecules, we trained the model for 5 epochs, whereas the largest datasets were trained for 2 epochs. Two accuracy measures were defined: (a) a triplet-based accuracy quantifying the proportion of correctly classified triplets by Euclidean distance measurements in the embedding space (dropping out S_i_); and (b) an anchor-based accuracy measuring the correlation between the S_i_-dropped-out embedding and the S_i_-only embedding. Given the bimodal distribution endowed by the Tanh activation, we chose to use a Matthews correlation coefficient (MCC) on a contingency table of binarized data (positive/negative along the 128 dimensions).

#### Light-weight signaturizers

We ran predictions for all molecules available in the CC universe (N = 778,531), producing 25 matrices of shape (n, 128). These matrices were used to learn chemistry-to-signature (CTS) signaturizers that are easy to distribute, allowing us to obtain signatures for a given molecule on-the-fly. CTS signaturizers were trained on a large number of molecules (N) with the aim to approximate the pre-calculated signatures presented in this work. Thus, in practice, a CTS signaturizer will often act as a mapping function, since the number of pre-calculated signatures is very large and covers a considerable portion of the medicinal chemistry space. CTS signaturizers were trained for 30 epochs and validated with an 80:20 train-test split, using 2048-bit Morgan Fingerprints (radius = 2) as feature vectors. Three dense hidden layers were used (1024, 512 and 256 dimensions) with ReLU activations and dropout regularization (0.2). The output was a dense layer of 128 dimensions (Tanh activation). The Adam optimizer was used (learning rate = 10^−3^). CTS signaturizers achieved a correlation with the type III signature of 0.769 +/− 0.074.

### Applicability domain estimation

An applicability score (α) for the signatures can be obtained at prediction time by means of a linear combination of five factors related to three characteristics that help increase trust in the predictions. These factors were tuned and calibrated on the test set.

#### Distance

Signatures that are close to training-set signatures are, in principle, closer to the applicability domain. We measured this distance in an unsupervised way (i.e. average distance to 5/25 nearest-neighbors) and in a supervised way by means of a random forest regressor trained on signatures as features and prediction accuracy (correlation) as dependent variable. In addition, we devised a measure of ‘intensity’, defined as the mean absolute deviation of the signatures to the average (null) signature observed in the training set.

#### Robustness

The signature-dropout procedure presented above can be applied at prediction time to obtain an estimate of the robustness of the prediction. For each molecule, we generated 10 dropped-out inputs, thereby obtaining an ensemble of predictions. Small standard deviations over these predictions indicate a robust output.

#### Expectancy a priori

We calculated the accuracy that is expected given the input signatures available for a particular molecule. Some CC signature types are highly predictive for others; thus, having these informative signatures at hand will in principle favor reliable predictions. This prior expectancy was calculated by fitting a random forest classifier with 25 absence/presence features as covariates and prediction accuracy as outcome.

### Library enrichment for activity against Snail1

#### Computational screening

##### Compound collections

Two compound collections were considered for screening, namely the IRB Barcelona library (17,563 compounds, considering the connectivity layer of the InChIKey) and the commercial Prestwick library (1,108 compounds). Of these, 627 and 1,104 were part of the CC universe, respectively, meaning that they had some type of reported bioactivity.

##### Chemical queries

These queries involved the search for compounds that were (i) chemically similar to curated DUB inhibitors, based on their known activity on promising DUBs according to a previous siRNA/Snail1 screen^18^ (Data S1), or (ii) similar to DUB inhibitors belonging to a broader list (with DUB-binding data available). Query i was achieved by computing chemical similarity (best across A1-2, P < 0.001) to a DUB inhibitors from the literature (curation categories 1 and 2 in Data S1, corresponding to 6 DUB inhibitors); 56 compounds were selected by this query. Query ii was composed of the intersection between two sub-queries. First, we looked for compounds that were similar (A1-2, P < 0.001) to any DUB inhibitor from the literature (all curation categories in Data S1). Of these, with a specific known (or predicted) compound-DUB interaction, according to a high-confidence binding collection extracted from B1, B2 and B4 (below the Pharos cutoff^42^). Compound-DUB interactions were predicted by using an in-house version of the classical similarity ensemble approach (SEA)^43^ based on A1-2, and taking into account ‘maximum similarity’, as recently recommended by SEA authors^44^ (we chose the cutoff with an optimal precision-recall trade-off).

##### Biological queries

In addition to DUBs, we considered other proteins relevant to Snail1 activity, namely TGFBR1/2, ERK2, FBXL5/14, DDR2 and GSK3B (Data S1). We then looked for transcriptional signatures associated with these genes in the L1000 Connectivity Map (shRNA assays, reversed over-expression assays, and known small molecule perturbagens) and also in CREEDS, which brings together data from GEO^45^. Overall, we gathered 132 transcriptional signatures with potential of having a connection to Snail1 (Data S1). Different priorities (0-4) were given to these signatures based on our mechanistic knowledge of Snail1 (Data S1 legend). Transcriptional signatures were converted to the CC D1 format as explained above. In addition, we derived C3-5 signatures for the Snail1-related genes, including DUBs highlighted by the siRNA/Snail1 screen.

We looked for connectivities (similarities, P < 0.001) between signatures of compounds in the D1 space and the list of Snai1-related signatures (at least 10 up/down regulated genes per signature). We did two searches (search H and search L), one against high-priority signatures (priority ≥ 3, Data S3), and another with a more relaxed cutoff (priority ≥ 1). In parallel, we derived C3-5 signatures for non-DUB Snail1-related genes (TGFBR1/2, etc.).

##### Random query

Molecules were randomly picked from the PWCK and IRB libraries, proportionally to the relative abundance of molecules from the two libraries in the lists retrieved from the previous queries (Data S1).

#### Cells

We used MDA-MB-231 cells stably transduced with pLEX-Snail1-Firefly Luciferase and pMSCV-Renilla Luciferase from our previous study^18^, and cultured them in DMEM supplemented with 10% FBS, glutamine and antibiotics (Thermo Fisher Scientific).

#### Dual-luciferase assay screening

We seeded 5·10^4^ cells in 96-well white plates prepared for cell culture (Corning). The day after, pre-diluted compounds of the chemical libraries were added to the cells at a final concentration of 20 μM, or in a few cases, of 4 μM, depending on the stock concentration and the maximum amount of DMSO that could be used in the assay. Several replicas of the vehicle controls (DMSO) or the positive control (the general DUB inhibitor PR-619 (Sigma-Aldrich)) were distributed along the experimental plates to allow internal normalization. After 6 h of incubation, medium was removed. Cells were then directly lysed with passive lysis buffer (Promega), and plates were stored at −20°C. Firefly and Renilla luciferase were quantified using the Dual-Luciferase Reporter assay system (Promega) in a GloMax luciferase plate reader (Promega). Four replicas conducted on two days were performed.

Intensities were corrected for each measurement (i.e. Firefly and Renilla) using one linear model per replica. The linear model included plate, row and column (as ordinal covariates) and type of measure (namely compounds, negative and positive controls) as fixed effects, as well as plate-row and plate-column interactions. Estimation of effects for plate, row and column (and their interactions) were used to correct intensity values. Intensities were previously transformed (square root) in order to fulfill the assumptions of linear models. In practice, this transformation implies a correction based on the median (instead of mean) effects, and it is thus robust to outliers (potential hits). Corrected values were transformed back to the original scale of the measures after correction. For normalization against controls, log2-ratios of intensities were computed against the mean of negative controls within each marker-replicate. Log2-ratios of Firefly:Renilla were then computed for signal evaluation.

The enrichment of hit rates was evaluated separately for each query (chemical, biological) with respect to the random distribution of Firefly:Renilla ratios.

### Signature-activity relationship (SigAR) models

For each classification task in the MoleculeNet, we sought to predict active/inactive (1/0) compounds using horizontally stacked CC signatures. A random forest classifier was trained using hyperparameters identified with HyperOpt^46^ over 10 iterations (number of estimators: (100, 500, 1000), max depth: (None, 5, 10), minimum sample split: (2, 3, 10), criterion: (gini, entropy), maximum features: (square root, log2)). Classifiers were calibrated using a Mondrian cross-conformal prediction scheme over 10 stratified splits. Evaluation was done with five stratified 80:20 train-test splits. Large MoleculeNet datasets such as PCBA were trained on a maximum of 30 under-sampled datasets, each comprising 10,000 samples. Scaffold-aware stratified splits, when necessary, were done ensuring that Murcko scaffolds^47^ observed in the training set were not present in the test set^48^.

Signature importance for each prediction was calculated by aggregating Shapley values (SHAP) as follows. First, features were ranked by their absolute SHAP across molecules. We then calculated the cumulative rank specific to each signature type (S_i_) (up to 250 features). Signature types with more of their dimensions in highly ranked positions were deemed to be more ‘explanatory’ for the prediction task.

## Acknowledgements

We would like to thank the SB&NB lab members for their support and helpful discussions. We are grateful to T.O. Botelho, I. Ramos and C. Gonzalez for giving us access to the IRB Barcelona and Prestwick libraries. P.A. acknowledges the support of the Generalitat de Catalunya (RIS3CAT Emergents CECH: 001-P-001682 and VEIS: 001-P-001647), the Spanish Ministerio de Economía y Competitividad (BIO2016-77038-R), the European Research Council (SysPharmAD: 614944) and the European Commission (RiPCoN: 101003633).

## Author contributions

M.D-F. and P.A. designed the study and wrote the manuscript. M.B., M.D-F., P.B-i-M., M.O-R. and O.G-P. implemented the entire computational strategy. EP, VA, VMD, AB-Ll and AGdH performed and analysed the Snail1 luciferase assays. All authors analyzed the results and read and approved the manuscript.

## Supplementary Figures and Tables

**Figure S1.**
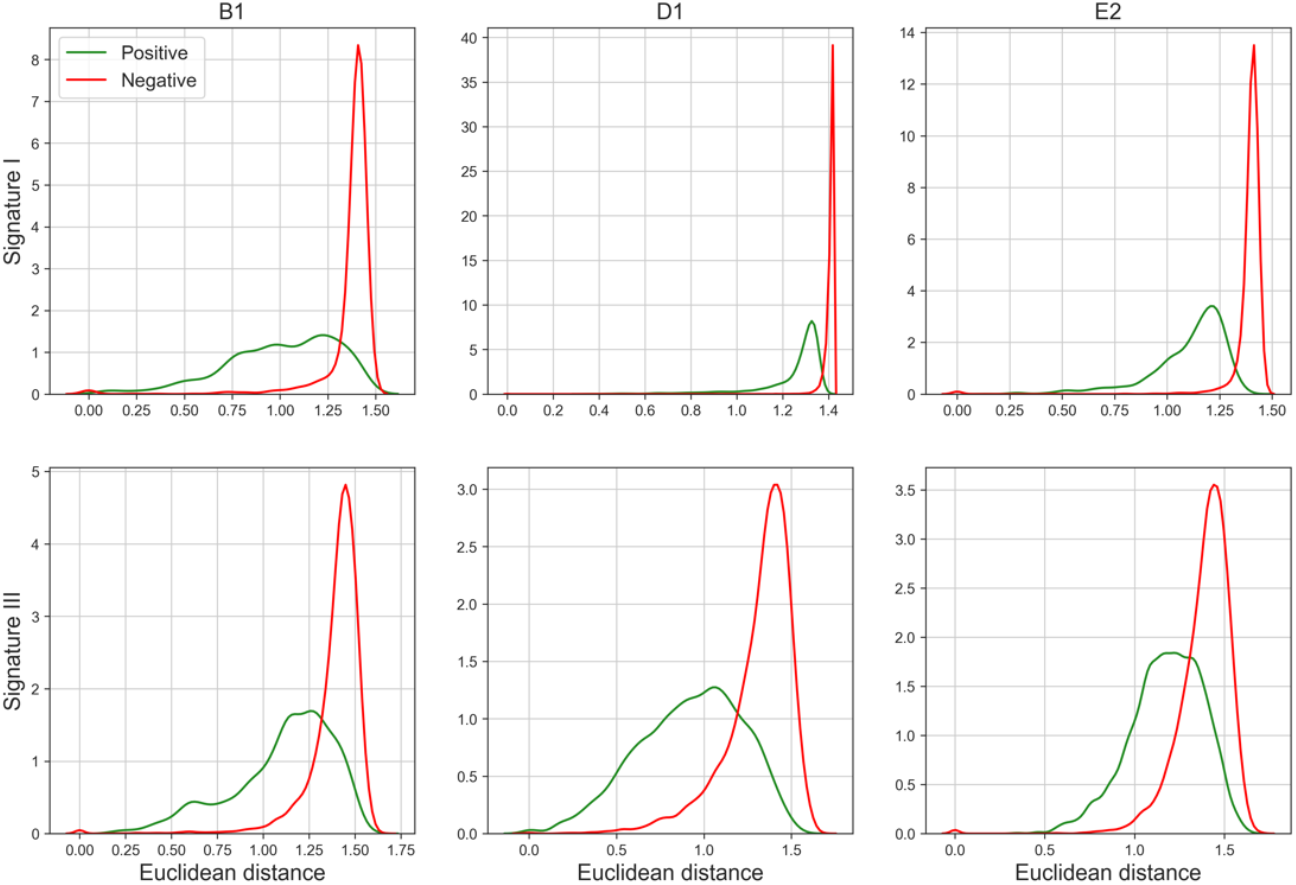
Original and learnt triplet distances for three representative CC datasets, namely B1, D1 and E2. The upper row shows the anchor-positive (green) and anchor-negative (red) Euclidean distances observed in the signature type I space (i.e. experimental signatures). Positive samples are closer to the anchor than negative ones. Correspondingly, the bottom row shows the distances observed in the signature III space (i.e. SNN embedding). Only test-test comparisons, where none of the molecules were seen during training, are shown.

**Figure S2.**
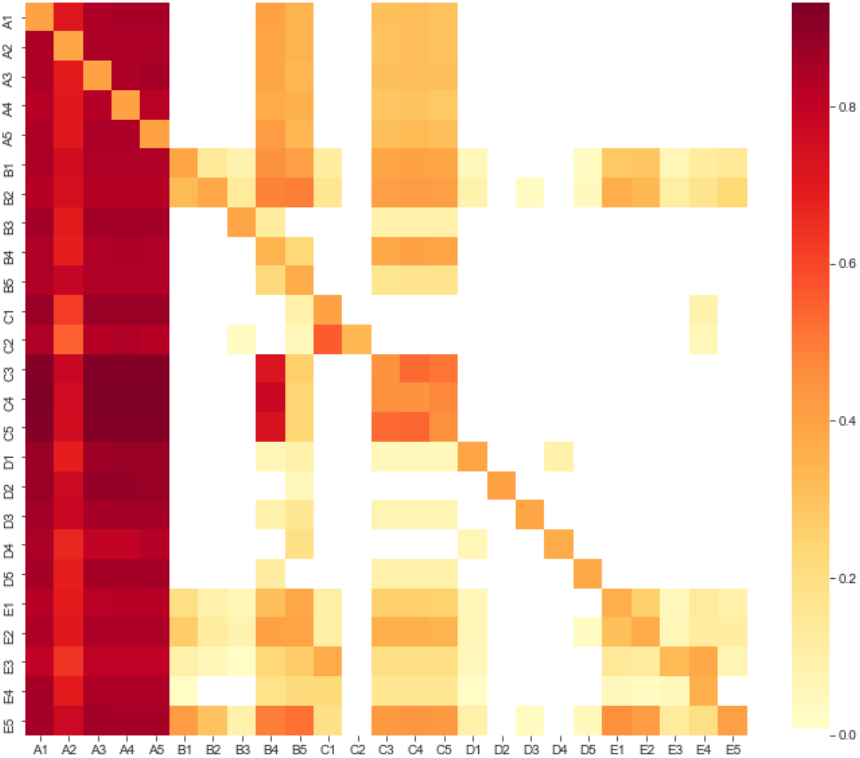
Proportion of signatures kept by the signature-dropout strategy. Rows (i) represent the CC space for which the SNN is being trained, and columns (j) correspond to the signatures being sampled. Red indicates that j signatures were typically used to train the i signaturizer.

**Figure S3.**
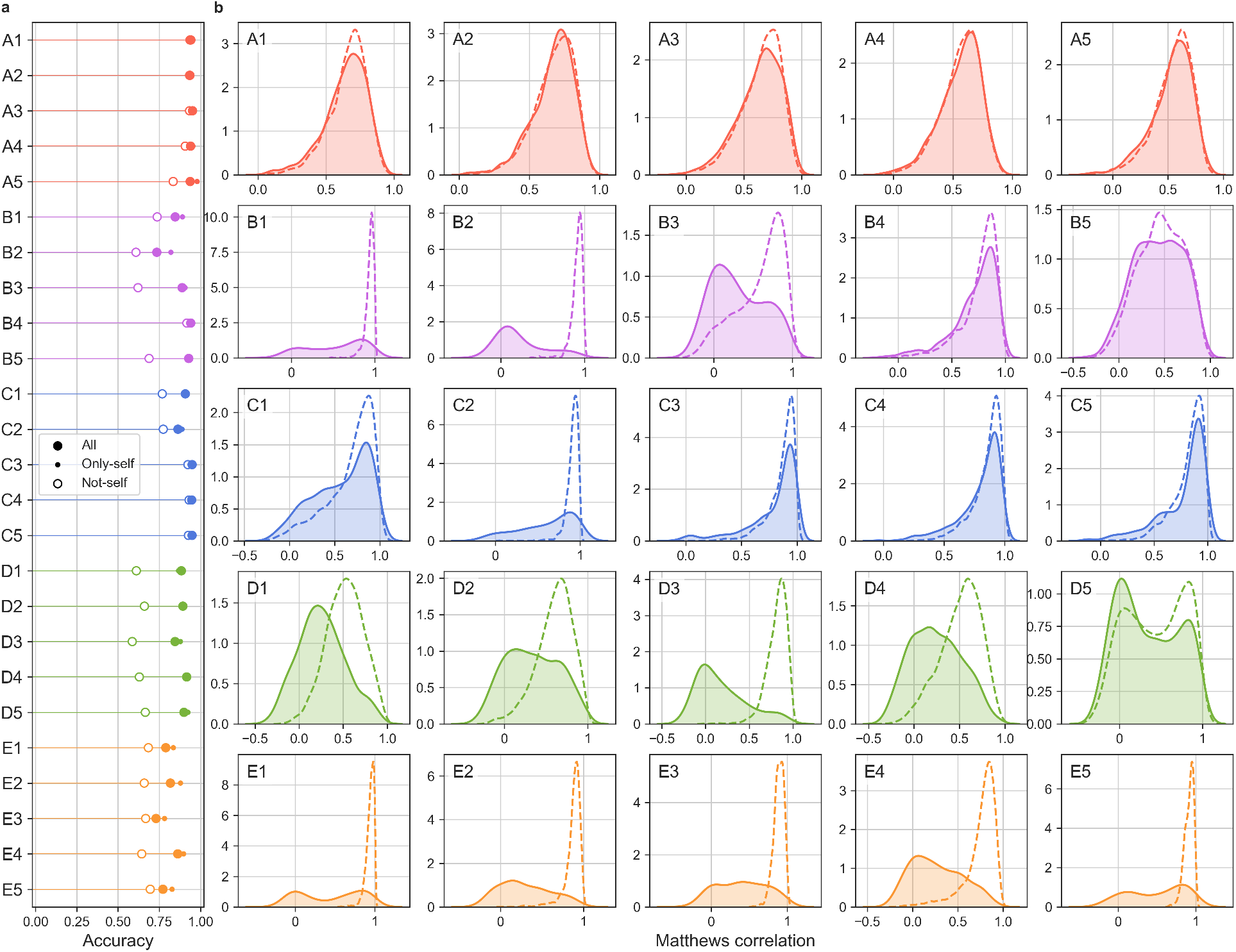
Performance of the signaturizers. (a) Performance (measured as a triplet-resolving accuracy) of signatures produced using all data (including the space of interest, ‘all’), only the space of interest (‘only-self’), and not using the space of interest (‘not-self’). Related to Figure 1C. (b) MCC scores (‘predicted’ signature vs ‘known’ signature) for train and test samples, depicted as dashed lines and filled shapes, respectively. Related to Figure 1d.

**Figure S4.**
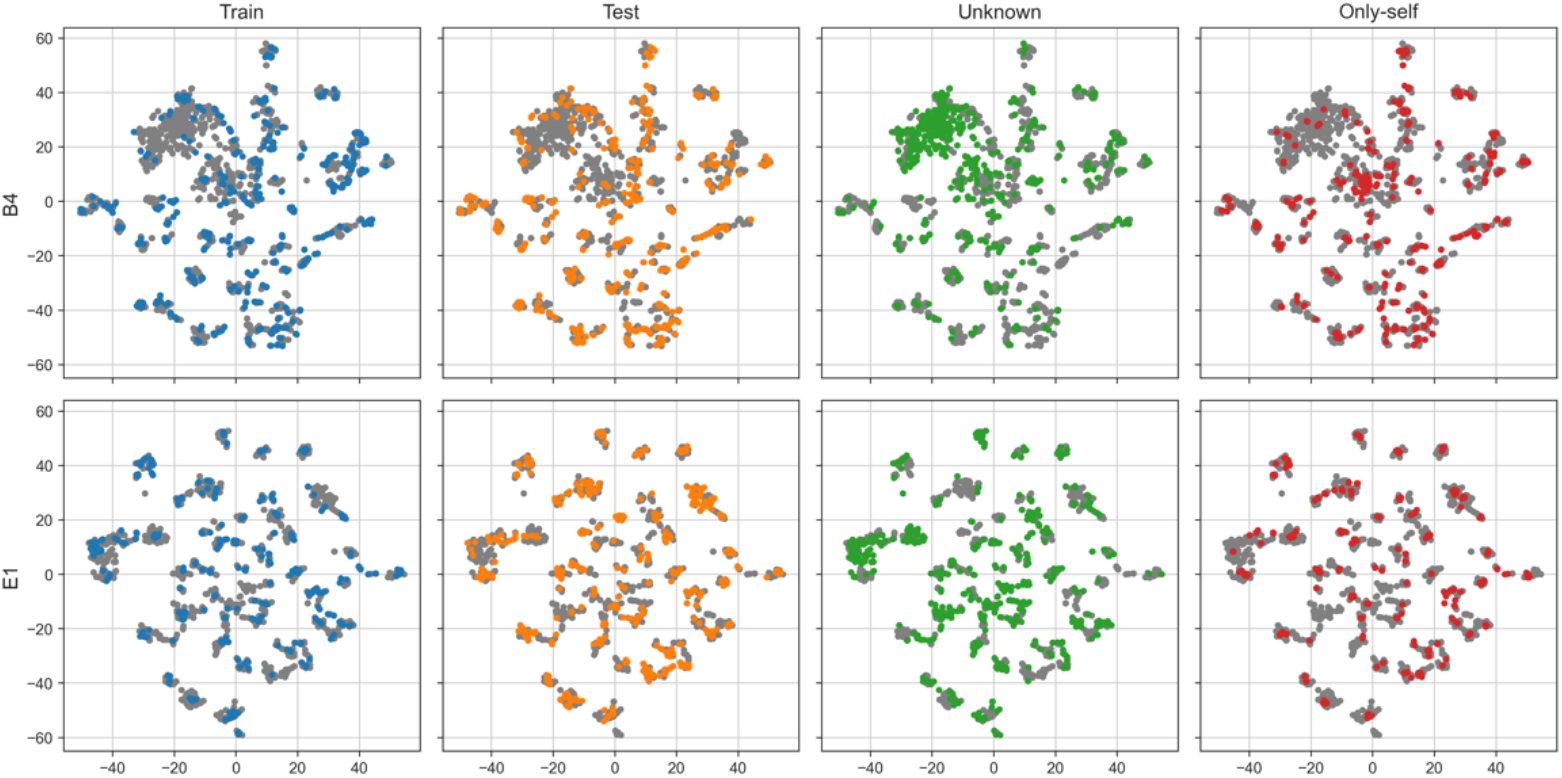
t-SNE 2D projections for two exemplary datasets (B4 and E1). The first two columns correspond to molecules in the training and test sets. ‘Unknown’ refers to signatures obtained for molecules with no available annotation in the space. ‘Only-self’ shows predictions done taking only the B4/E1 space as input.

**Figure S5.**
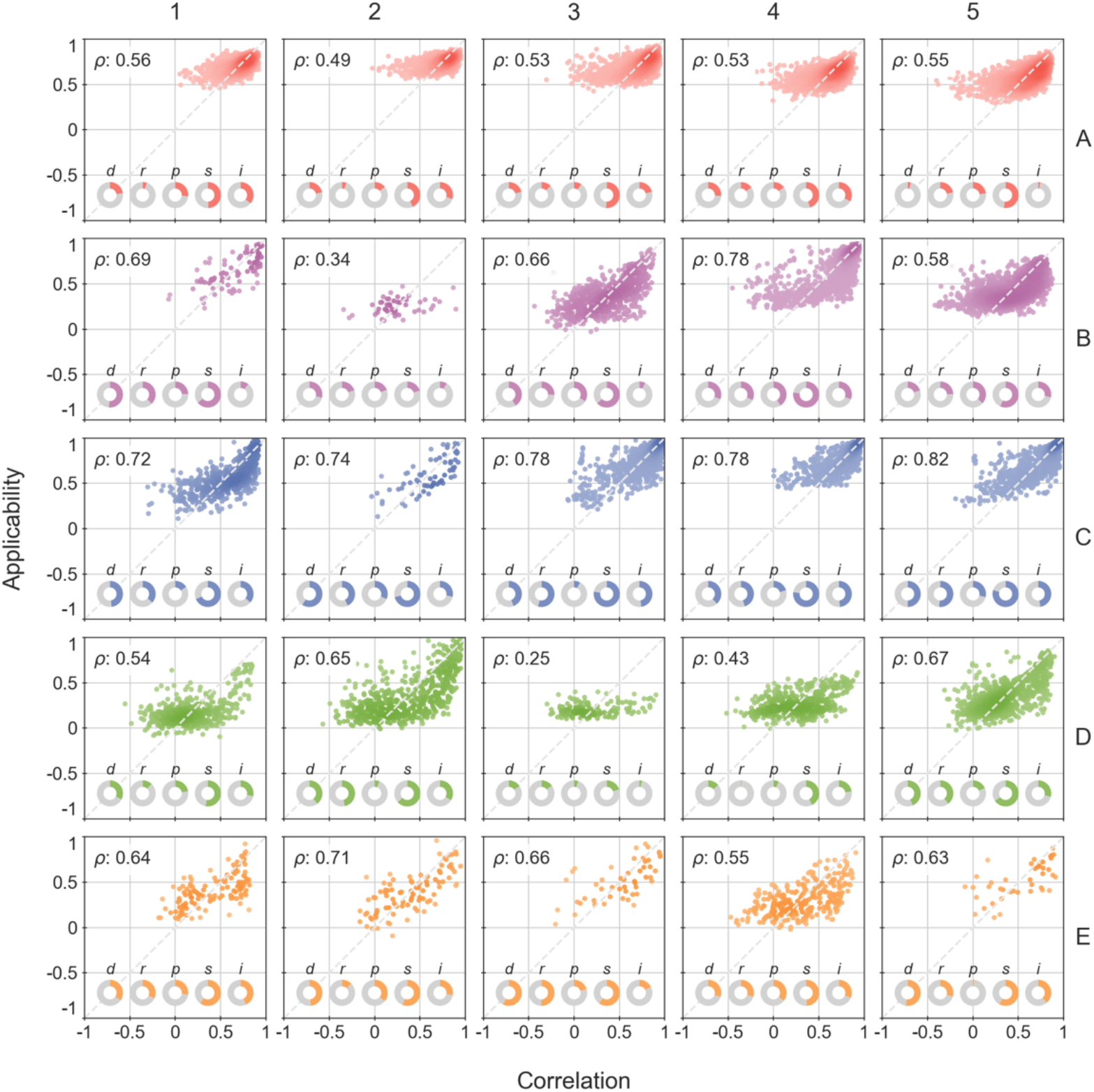
Correlation between the applicability score (α) and the true-vs-predicted signature correlation. The applicability score is determined by the linear combination of five factors, represented in the pie charts and abbreviated as follows; d: nearest-neighbor distance, r: robustness, p: prior (i.e. expected accuracy *a priori*), s: supervised distance, and i: intensity. The area covered by the pie chart corresponds to the coefficient of the linear combination to adjust an α score. Plots correspond to 80:20 train-test splits.

**Figure S6.**
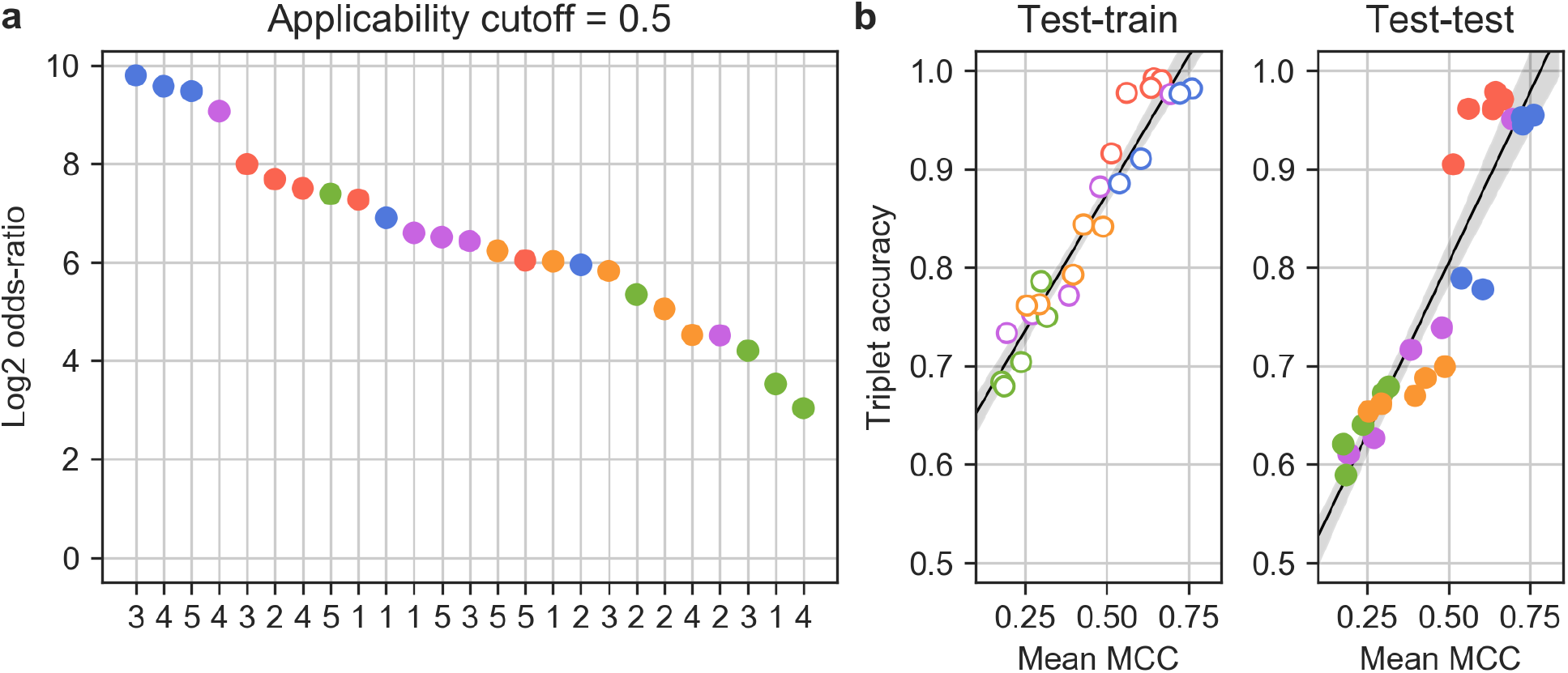
(a) Enrichment of GSig 10-nearest-neighbors at an α cutoff of 0.5, measured as the log_2_-odds ratio on a contingency table counting the number of neighbors common to S_i_ and GSig. High enrichments mean that, in the light of the global information available from the CC (i.e. GSig), similarities encountered for the S_i_ signature are relevant. The 25 CC categories are ranked by enrichment score. Color of the dot denotes CC level (A-E) and numbering indicates the sublevel (1-5). (b) Correlation between the two accuracy metrics (i.e. MCC and triplet accuracy) for train-train and test-test validations.

**Figure S7.**
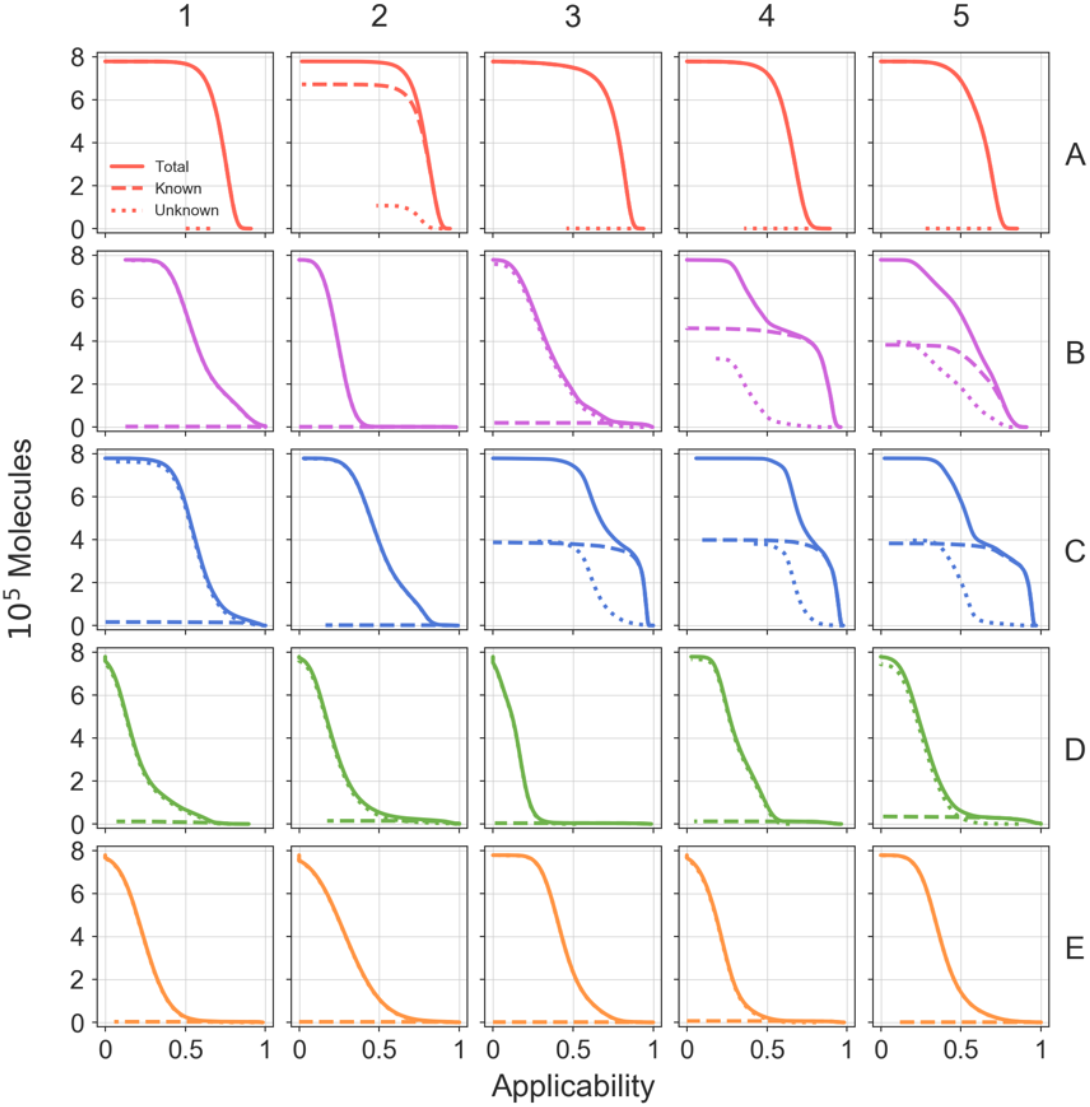
Number of molecules at different applicability (α) cutoffs, obtained for the full universe of CC molecules (~800k). The dashed line indicates the number of molecules with ‘experimental’ data in the corresponding CC space, while the dotted line indicates the number of molecules with no available data in the CC space. The solid line shows the total number of molecules. We can see that, for instance, with an α cutoff of 0.5 the number of molecules with reliable D1 signatures is 5-fold the number of molecules with experimental information.

**Figure S8.**
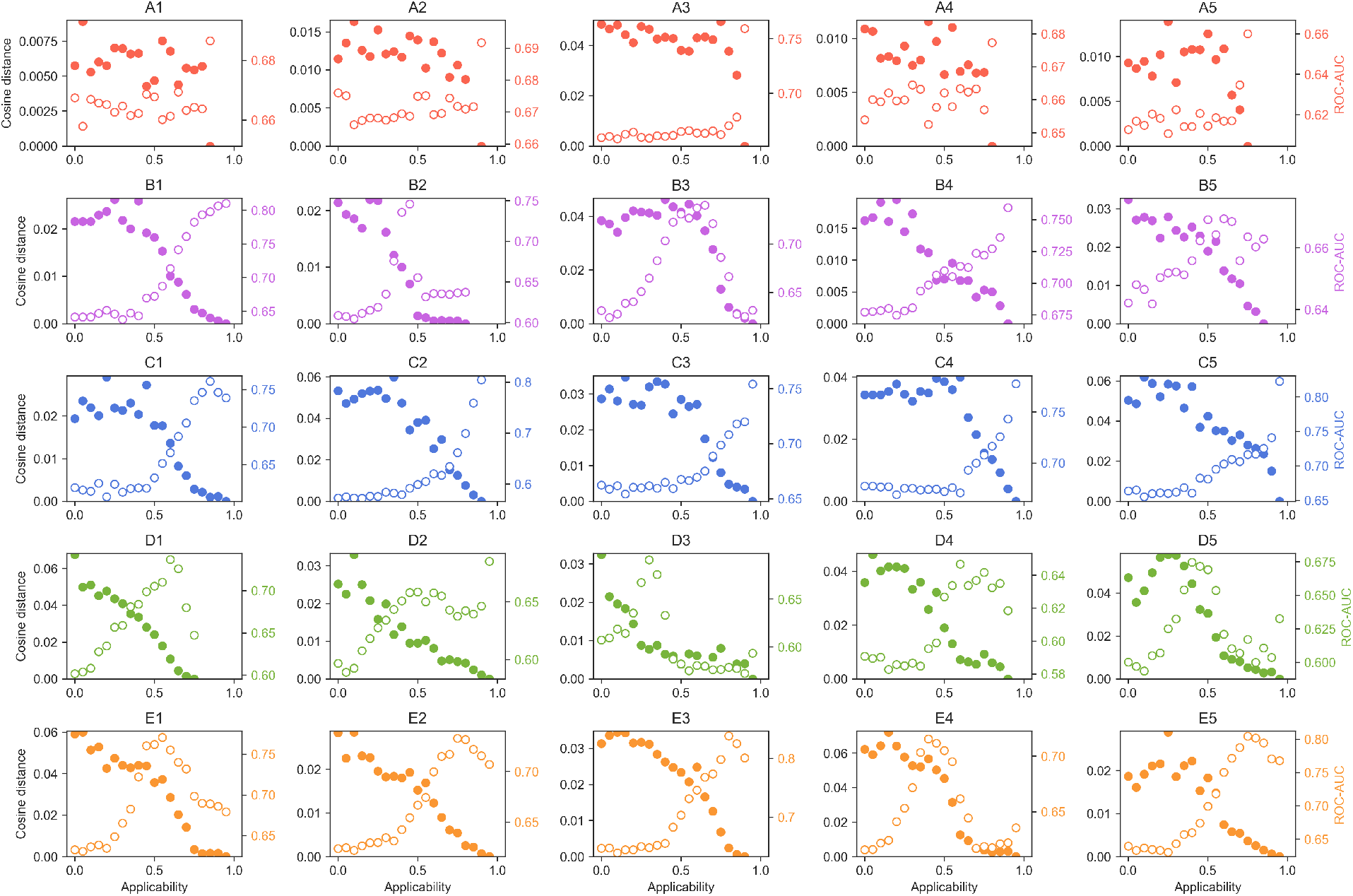
Exploration of the α score. For each CC dataset S_i_, we measure, at a given α cutoff, the capacity to recall 10-nearest-neighbors across CC spaces, similar to what is done in Figure 2b. The average ROC-AUC of S_i_ along S_1-25_ is plotted as empty dots (right, colored axis). We consider the profile of ROC-AUCs obtained at the highest α to be the most genuine for the signature. We measure how the profile of ROC-AUCs diverges (cosine distance of the 25-dimensional ROC-AUC vector) as lower α values are taken. Larger distances indicate less ‘purity’ in the signature-correlation profile (filled dots, left axis).

**Figure S9.**
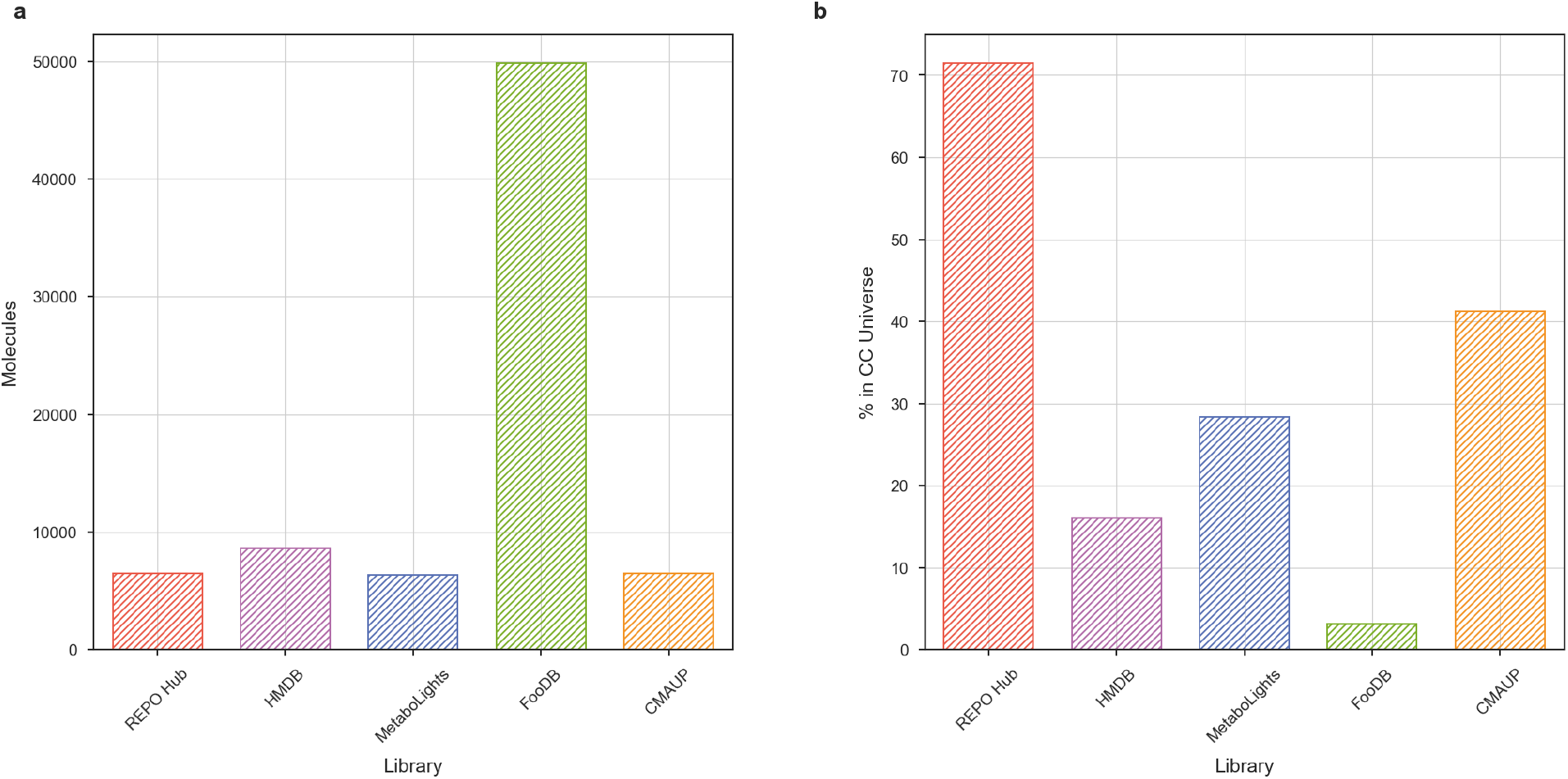
Number of molecules (a) and CC coverage (b) of five selected compound collections with respect to the CC universe (~800k molecules).

**Figure S10.**
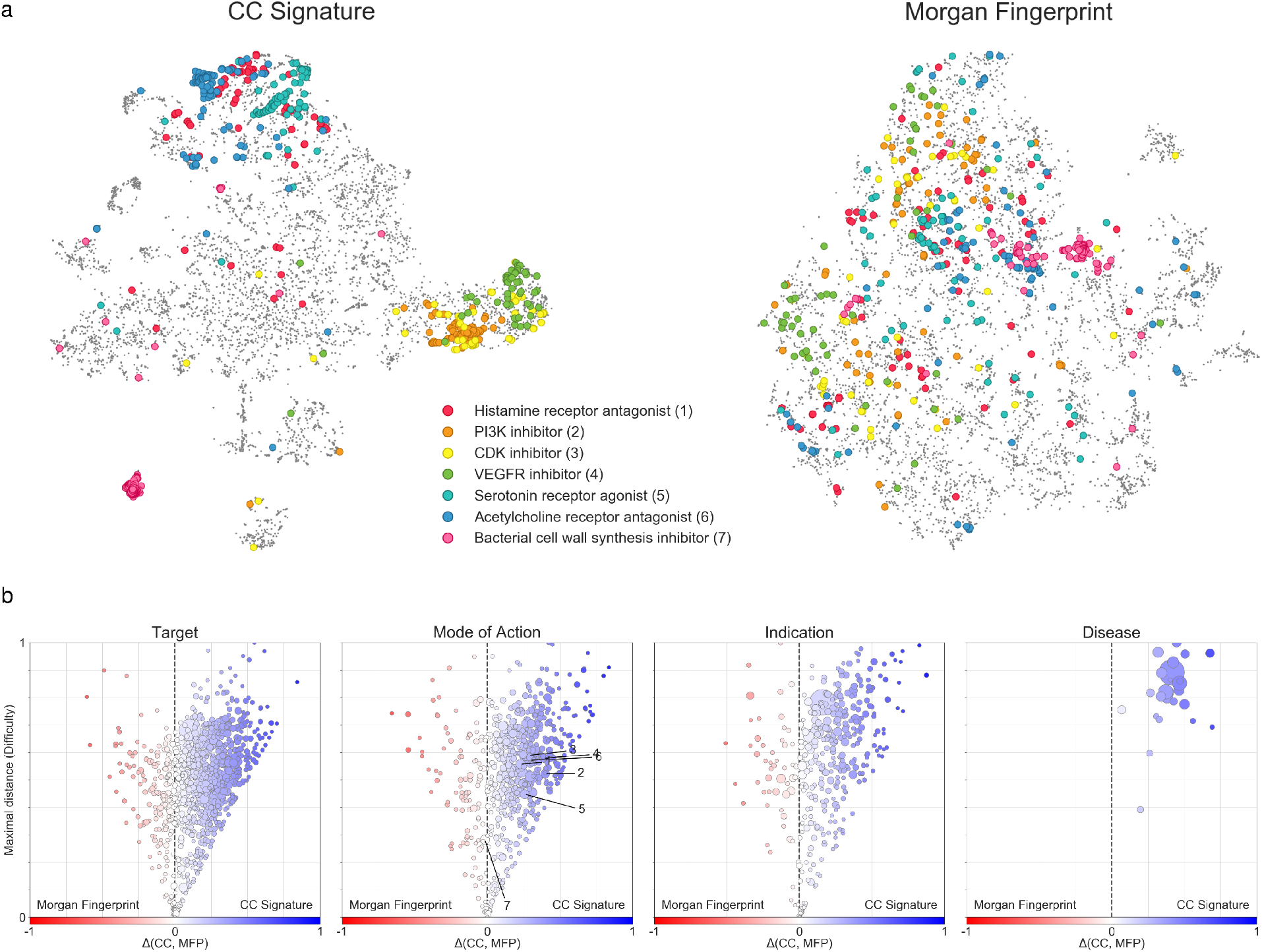
Drug Repurposing Hub 2D projections. (a) t-SNE 2D projections based on GSigs (left) compared to Morgan fingerprint (MFp; 2048-bit, radius: 2) projections (right). Regions corresponding to certain MoAs are highlighted. (b) Level of clustering of the different annotations specified in the Drug Repurposing Hub, namely ‘targets’, ‘MoA’, ‘indications’ and ‘disease areas’. Each dot corresponds to an annotation, and the size of the dot is proportional to the number of molecules. The average Euclidean distance in the 2D-projection between molecules with the same annotation is calculated, both for GSig- and MFp-based projections. For each annotation size, 100 randomly sampled points are drawn from the projection in order to scale the average distance measure. The x-axis measures the difference between GSig and MFp distances. Values close to 1 indicate that molecules of a certain annotation are well localized in the GSig projection and scattered in the MFp projection. Values close to −1 indicate the contrary. The red-to-blue color scale follows this axis. The y-axis is a measure of ‘difficulty’, i.e. the maximal inter-annotation distance observed between the GSig and MFp projections. Values close to one indicate that, in one of the projections the distance between molecules in the annotation is large (i.e. scattered points), while values close to 0 indicate that in both projections molecules with the same annotation are close-by. Thus, points in the upper-right corner are favorable to the CC projection, points in the mid-bottom region are well-grouped in both projections, and points in the upper-left corner are favorable to the MFp projection. The numbering in the MoA subplot relates to the legend in the top panel.

**Figure S11.**
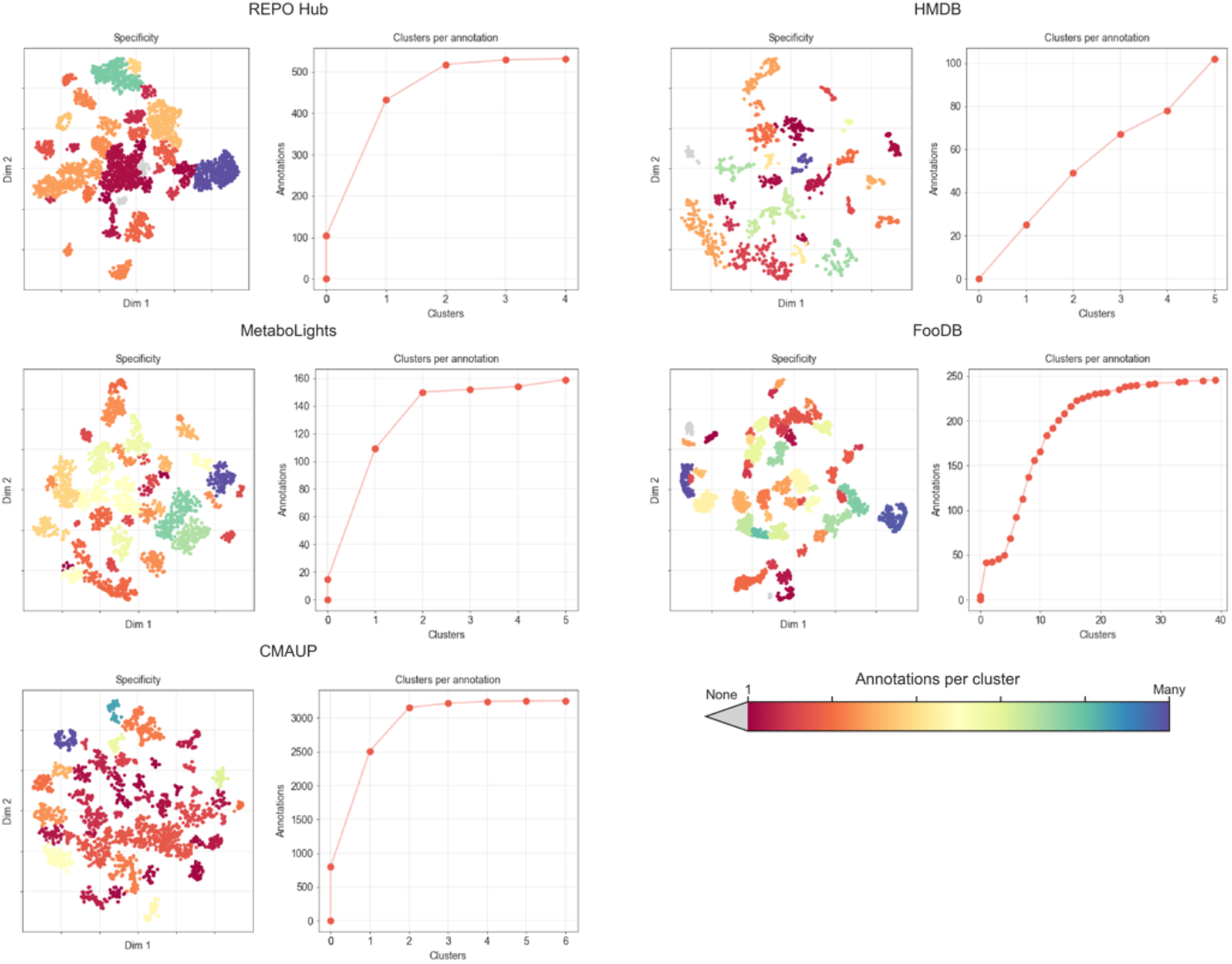
Association analysis between clusters discovered in 2D projections and annotations available for the compounds. REPO Hub, HMDB, MetaboLights, FooDB and CMAUP datasets are analyzed separately. REPO Hub annotations: targets, MoAs, indications, disease areas. HMDB annotations: tissues, biofluids, biofunctions, cellular components and origin. MetaboLights annotations: organism name, group, genus, species, etc. FooDB: food group, subgroup and name. CMAUP: plant family and species. The DBSCAN clustering algorithm was used to identify clusters based on the *(x,y)*-coordinates of the 2D projection. Then, a Fisher’s exact test was performed for each cluster-annotation pair, based on a contingency table counting the number of molecules in/out of the cluster and the number of molecules with/without annotations; P-values < 0.01 with an FDR < 0.1 and a log2 odds-ratio > 1.5 were considered to be significant. In the plots, the color of the projections denotes the number of annotations found to be statistically associated with each of the clusters. Blue regions are ‘unspecific’, in that they contain molecules belonging to multiple annotations. Red regions are more ‘specific’ as they are associated with few annotations. Gray clusters are not enriched with any annotation. The cumulative plots show the number of clusters associated with each annotation. In REPO Hub, for instance, most annotations are associated with one or two clusters, whereas most FooDB annotations are found enriched in five clusters.

**Figure S12.**
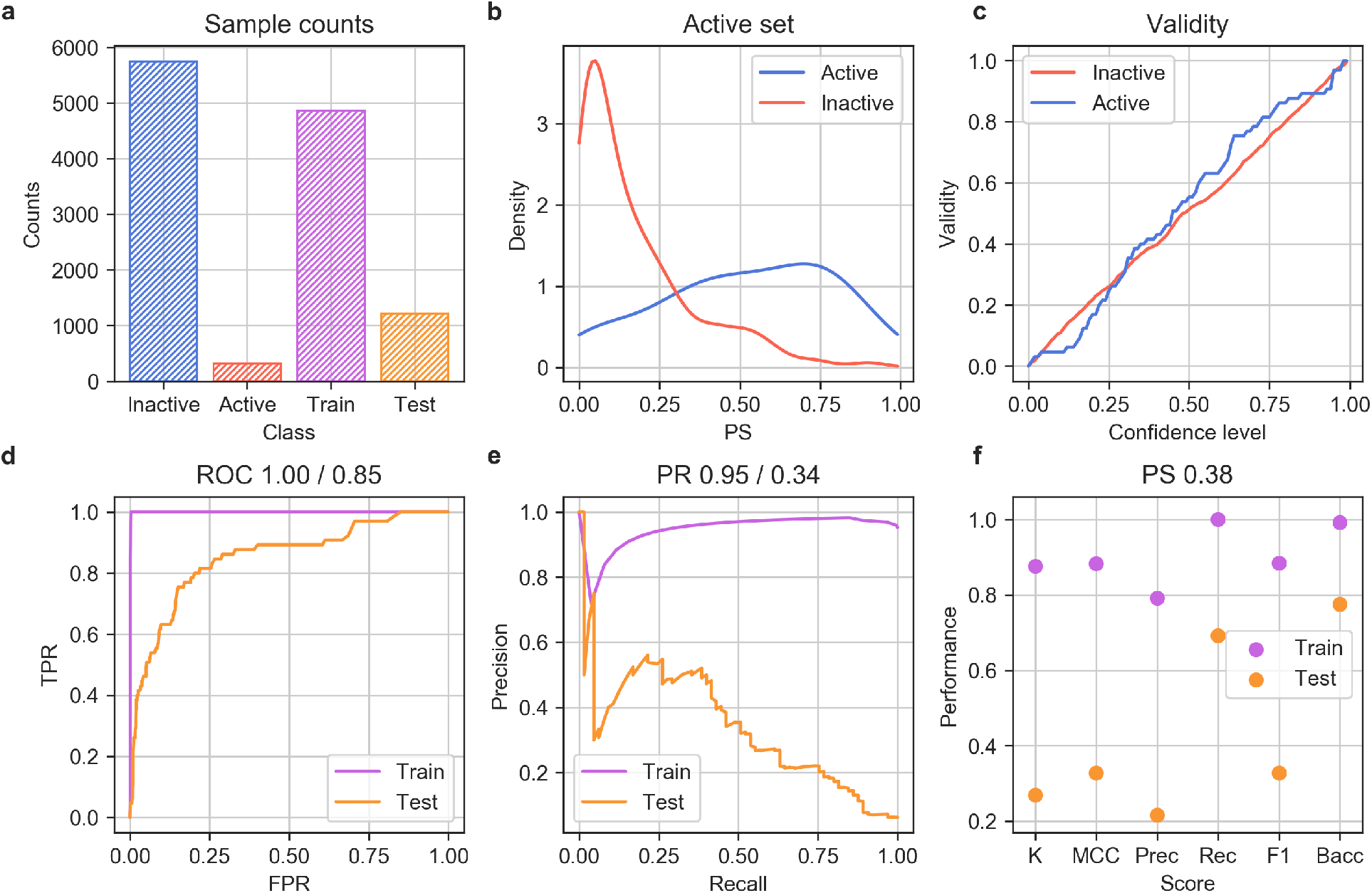
Evaluation of the SR-HSE random forest classifier. (a) Number of active and inactive molecules, and proportion of molecules in the train and test sets. (b) Prediction score (PS) assigned to active (blue) and inactive (red) molecules in the test set. (c) Validity plot of the cross-conformal predictor. (d) Train and test ROC curves. (e) Train and test precision-recall curves. (f) Other classification scores (PS > 0.38). K: Cohen’s kappa. Prec.: precision, Rec.: recall, Bacc: Balanced accuracy.

**Figure S13.**
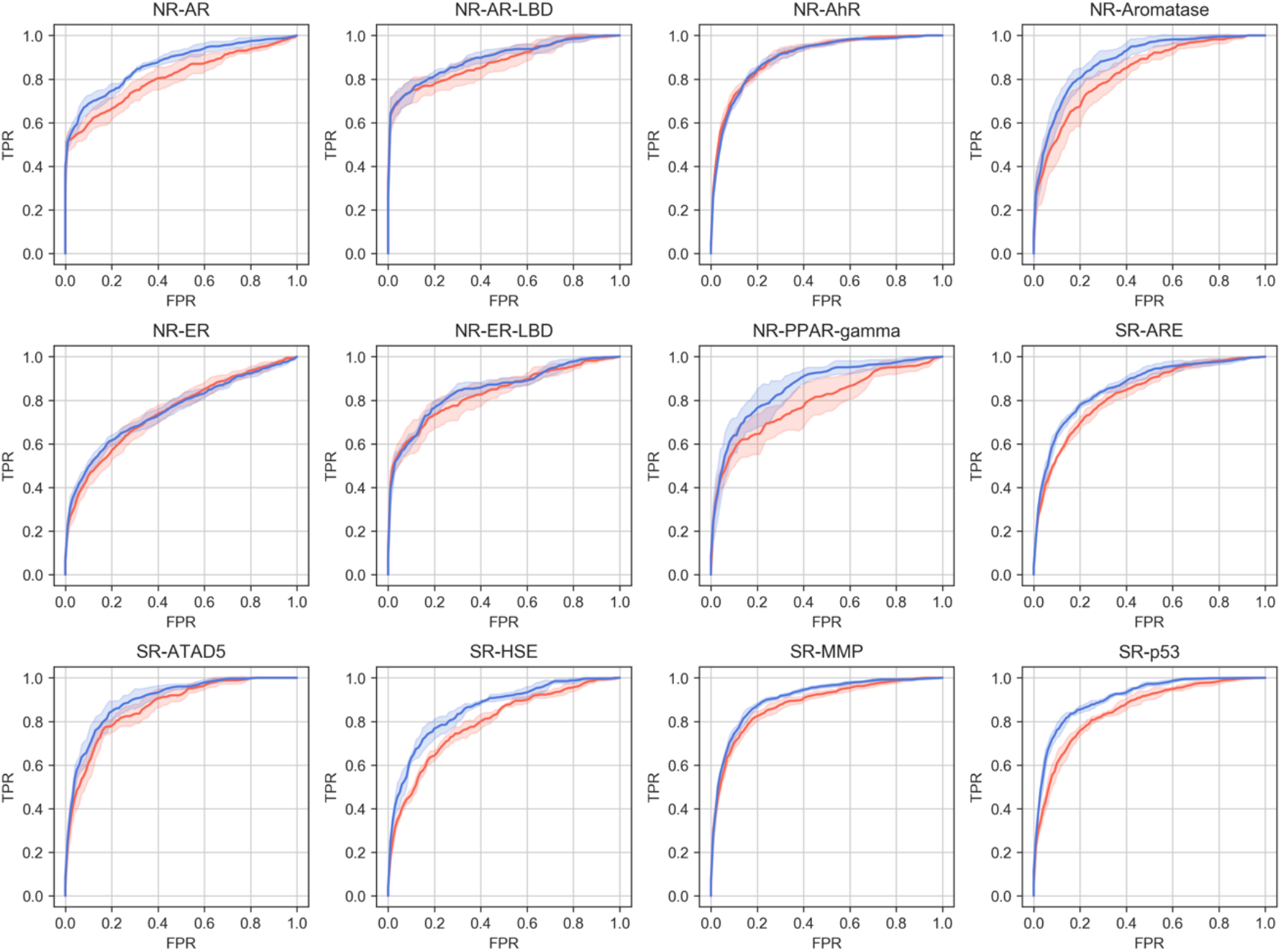
ROC curves for the 12 Tox21 prediction tasks. CC- and MFp-based predictors are blue and red, respectively.

**Figure S14.**
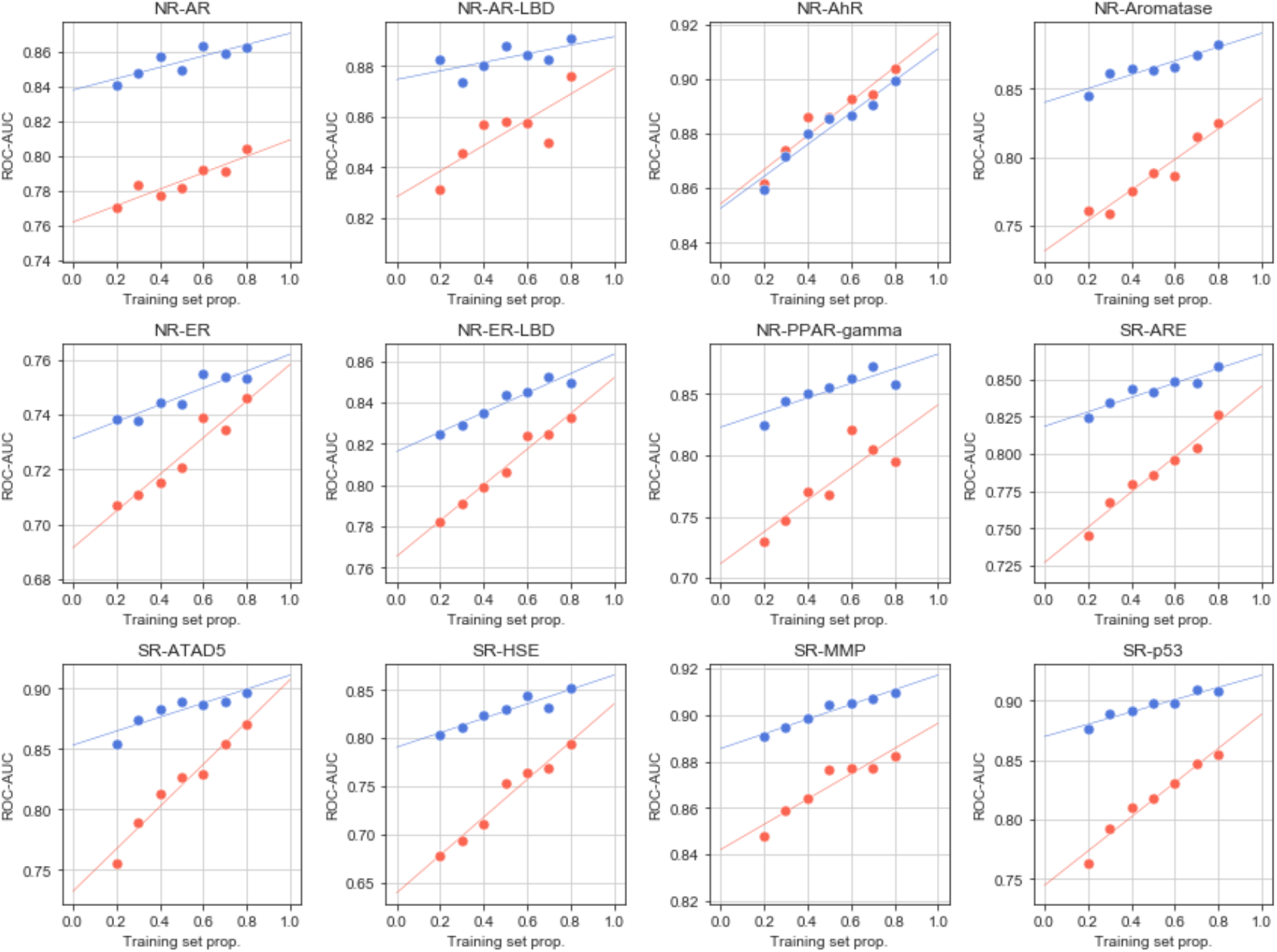
Performance (ROC-AUC) of the Tox21 models at different training set sizes and using GSigs (blue) and MFps (red) as feature vectors.

**Figure S15.**
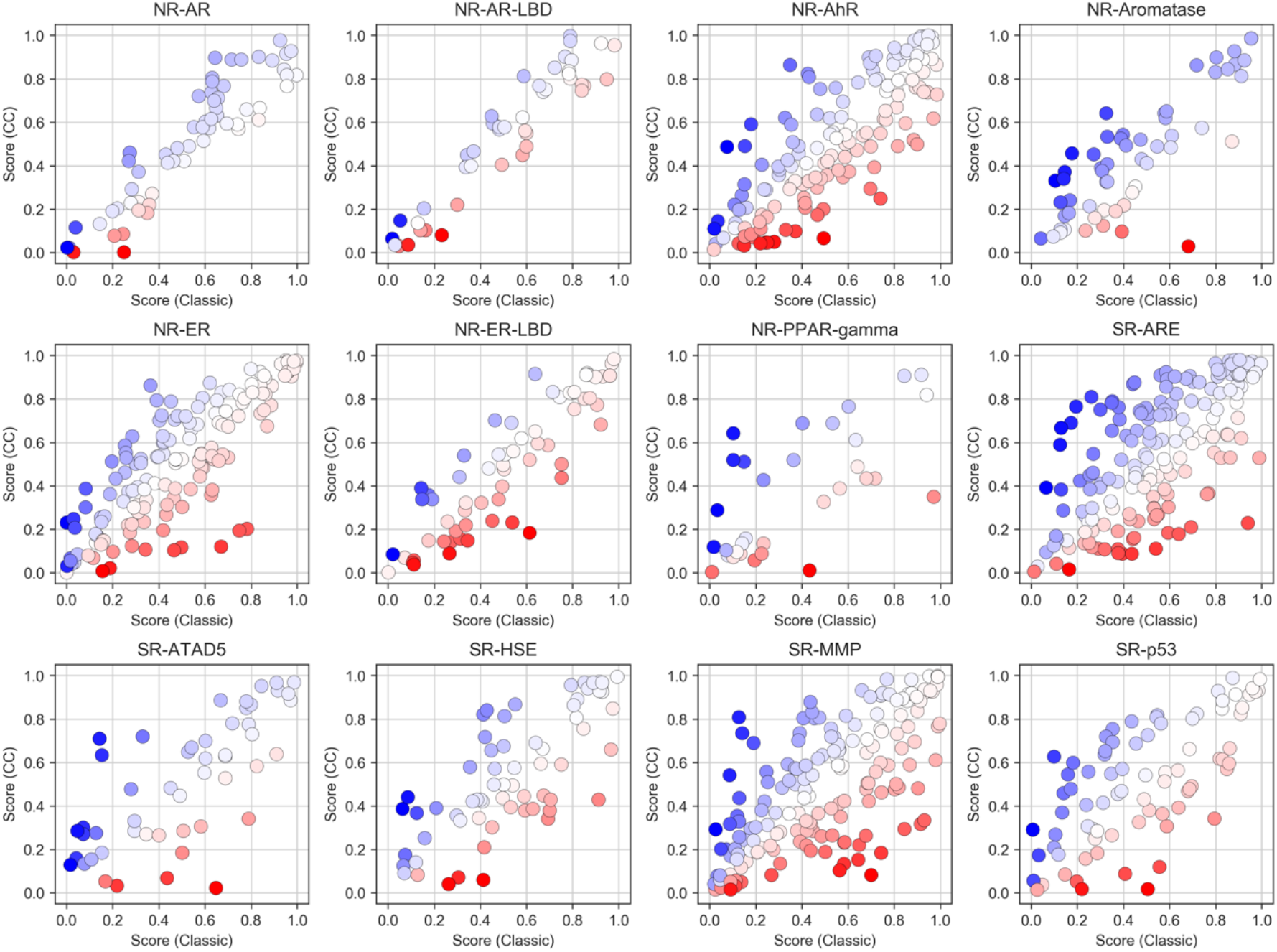
Prediction scores assigned to each active molecule by the predictors at test-time in a 5-fold cross-validation (Tox21 benchmark dataset). The color scale denotes the relative difference between CC scores and classic (MFp) scores. Blue indicates a high score by the CC predictor and a low score by the MFp predictor. Red indicates the opposite.

**Figure S16.**
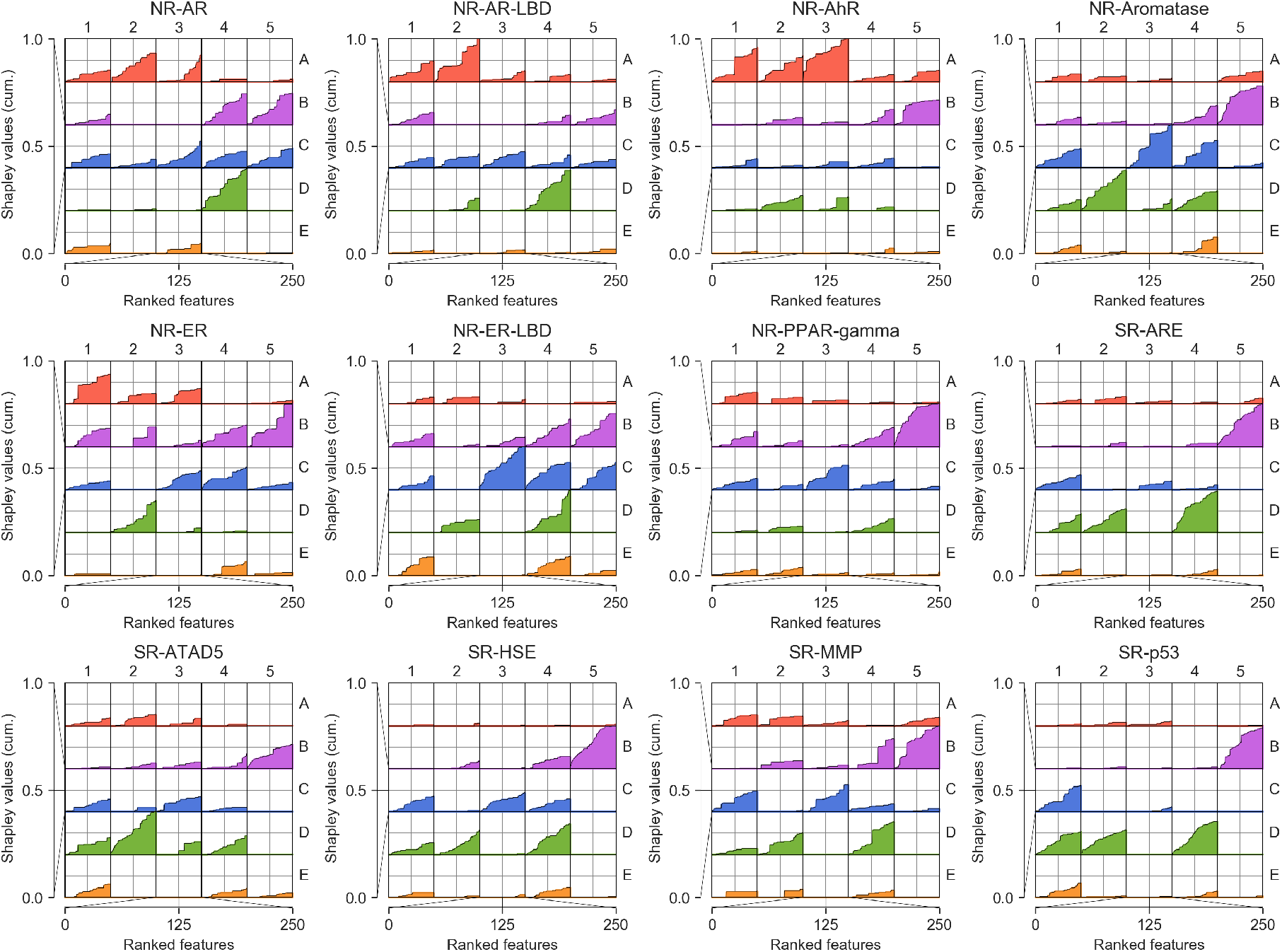
Explanatory potential of CC categories across the Tox21 benchmark dataset. Top 250 features (ranked by average absolute Shapley values across samples) are summed up in the corresponding S_1-25_ slots.

**Figure S17.**
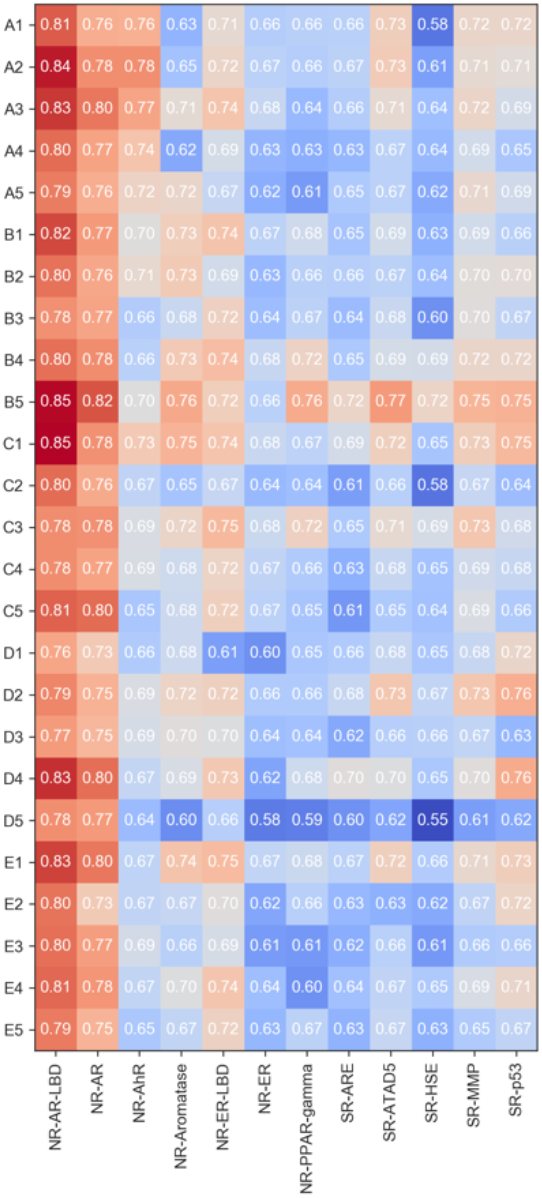
Discriminative power (ROC-AUC) of SVCs based exclusively on 2D representations of the signatures. The analysis is done for each signature type (A1-E5) and the 12 Tox21 tasks.

**Figure S18.**
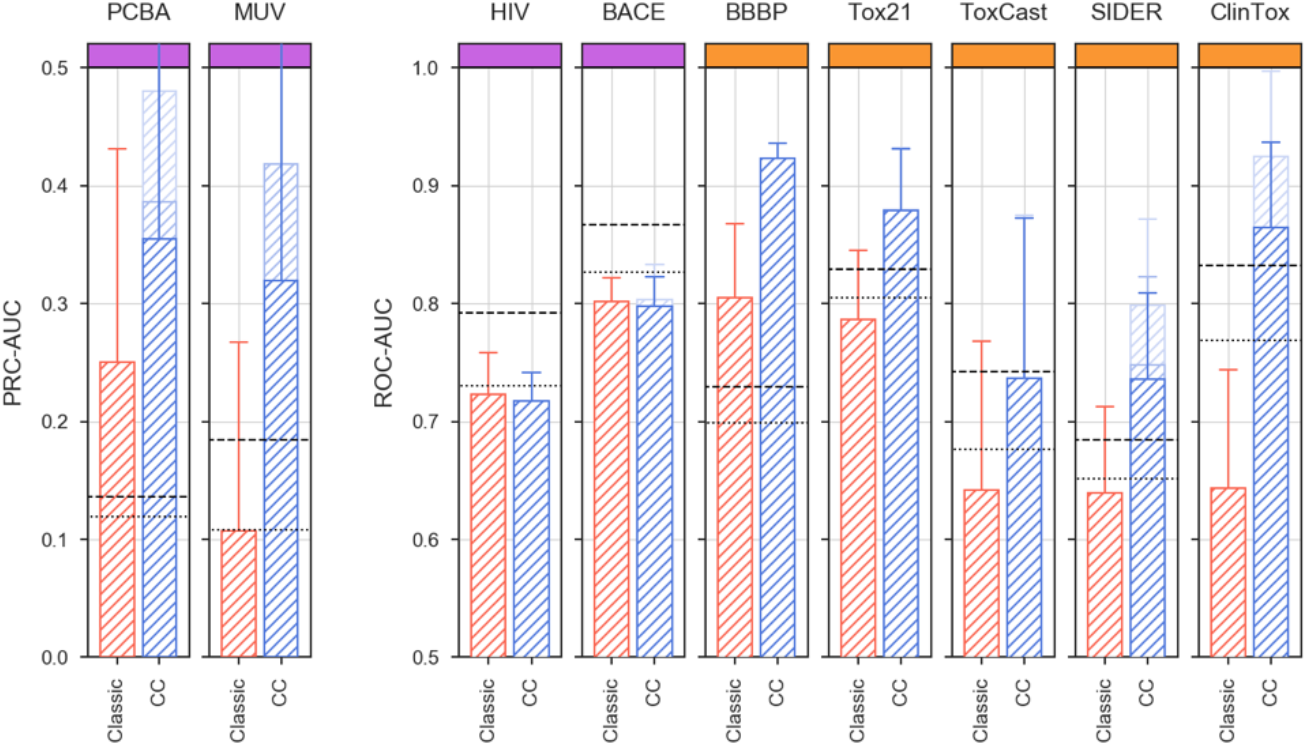
MoleculeNet validation done exclusively with molecules having at least one bioactivity data point previously available in the CC. The plot relates to Figure 4g (see legend for details).

**Table S1.** MoleculeNet classification tasks. ‘Removed’ column indicates the CC spaces that are not used to make predictions. The parenthesis denotes the most aggressive removal.

| *Category* | *Name* | *Molecules* | *Tasks* | *Split* | *Metric* | *Removed* |
| --- | --- | --- | --- | --- | --- | --- |
| Biophysics | PCBA | 437,929 | 128 | Random | PR-AUC | B5, (B4, C3, C4, C5) |
|  | MUV | 93,087 | 17 | Random | PR-AUC | B4, (B5, C3, C4, C5) |
|  | HIV | 41,127 | 1 | Scaffold | ROC-AUC | - |
|  | BACE | 1,513 | 1 | Scaffold | ROC-AUC | (B4, B5, C3, C4, C5) |
| Physiology | BBBP | 2,039 | 1 | Scaffold | ROC-AUC | - |
|  | Tox21 | 7,831 | 12 | Random | ROC-AUC | (E4) |
|  | ToxCast | 8,575 | 612 | Random | ROC-AUC | (E4) |
|  | SIDER | 1,427 | 27 | Random | ROC-AUC | E3, (E4) |
|  | ClinTox | 1,478 | 2 | Random | ROC-AUC | E1, E2, E3, E4, E5 |

**Data S1**. Library enrichment for activity against Snail1. This dataset contains the selected compounds from the IRB and PWCK libraries for their possible activity against Snail1, and the information we used to perform these queries (online Methods). We indicate and prioritize (1) known DUB inhibitors, listing their known targets, (2) DUBs according to the results of the siRNA-DUB/Snail1 screening assay and (3) public transcriptional signatures related with the Snail1 activation pathway. We report the score of each molecule in the inspected queries (e.g. similarity to known DUB inhibitors, reversion of Snail1 transcriptional signatures, belonging to the TGFβ pathway, etc.). We provide information for all compounds selected in chemical and biological queries, as well as the randomly selected ones.

